# Restoring hippocampal glucose metabolism rescues cognition across Alzheimer’s disease pathologies

**DOI:** 10.1101/2024.06.23.598940

**Authors:** Paras S. Minhas, Jeffrey R. Jones, Amira Latif-Hernandez, Yuki Sugiura, Aarooran S. Durairaj, Takeshi Uenaka, Qian Wang, Siddhita D. Mhatre, Ling Liu, Travis Conley, Hannah Ennerfelt, Yoo Jin Jung, Praveena Prasad, Brenita C. Jenkins, Ryan Goodman, Traci Newmeyer, Kelly Heard, Austin Kang, Edward N. Wilson, Erik M. Ullian, Geidy E. Serrano, Thomas G. Beach, Joshua D. Rabinowitz, Marius Wernig, Makoto Suematsu, Frank M. Longo, Melanie R. McReynolds, Fred H. Gage, Katrin I. Andreasson

**Author notes:** These authors contributed equally (Paras S. Minhas, Jeffrey R. Jones, Amira Latif-Hernandez).

## Abstract

Impaired cerebral glucose metabolism is a pathologic feature of Alzheimer Disease (AD), and recent proteomic studies highlight a disruption of glial carbohydrate metabolism with disease progression. Here, we report that inhibition of indoleamine-2,3-dioxygenase 1 (IDO1), which metabolizes tryptophan to kynurenine (KYN) in the first step of the kynurenine pathway, rescues hippocampal memory function and plasticity in preclinical models of amyloid and tau pathology by restoring astrocytic metabolic support of neurons. Activation of IDO1 in astrocytes by amyloid-beta_42_ and tau oligomers, two major pathological effectors in AD, increases KYN and suppresses glycolysis in an AhR-dependent manner. Conversely, pharmacological IDO1 inhibition restores glycolysis and lactate production. In amyloid-producing *APP^Swe^-PS1^ΔE9^* and 5XFAD mice and in tau-producing P301S mice, IDO1 inhibition restores spatial memory and improves hippocampal glucose metabolism by metabolomic and MALDI-MS analyses. IDO1 blockade also rescues hippocampal long-term potentiation (LTP) in a monocarboxylate transporter (MCT)-dependent manner, suggesting that IDO1 activity disrupts astrocytic metabolic support of neurons. Indeed, in vitro mass-labeling of human astrocytes demonstrates that IDO1 regulates astrocyte generation of lactate that is then taken up by human neurons. In co-cultures of astrocytes and neurons derived from AD subjects, deficient astrocyte lactate transfer to neurons was corrected by IDO1 inhibition, resulting in improved neuronal glucose metabolism. Thus, IDO1 activity disrupts astrocytic metabolic support of neurons across both amyloid and tau pathologies and in a model of AD iPSC-derived neurons. These findings also suggest that IDO1 inhibitors developed for adjunctive therapy in cancer could be repurposed for treatment of amyloid- and tau-mediated neurodegenerative diseases.

---

Alzheimer’s disease (AD) is an age-associated neurodegenerative disorder characterized by the progressive and irreversible loss of synapses and neural circuitry (*1*). Its prevalence is expected to triple by 2050 (*2*). Major pathophysiologic processes contributing to synaptic loss, including disrupted proteostasis, accumulation of misfolded amyloid and tau, and microglial dysfunction, are being vigorously investigated with the goal of identifying disease-modifying therapies. However, coincident with these distinct pathologies is a sustained decline in cerebral glucose metabolism (*3, 4*), with recent proteomics revealing a marked disruption of astrocytic and microglial metabolism in AD subjects (*5*). Astrocytes, which exist in approximately a 1:1 ratio to neurons (*6*), are essential for regulation of neurotransmitter levels and bioenergetic support of neurons. Specifically, astrocytes generate lactate that is exported to neurons to fuel mitochondrial respiration and support synaptic activity (*7–9*).

Recent studies have suggested a role for IDO1, an enzyme expressed in astrocytes that metabolizes tryptophan to KYN, in multiple neurodegenerative disorders, including AD (*10*). IDO1 is the rate limiting enzyme in the conversion of tryptophan to KYN, a metabolite that elicits immune suppression in inflammatory and neoplastic contexts via interaction with the aryl-hydrocarbon receptor (AhR) (*11*). In the brain, IDO1 is expressed in astrocytes and microglia but not in neurons, and levels increase in response to inflammatory stimuli (*12, 13*). Here we report that KYN generated by IDO1 suppresses astrocytic lactate transfer to neurons in mouse models of AD pathology and human iPSC-derived astrocytes from late-onset AD subjects. Genetic deletion of IDO1 or pharmacologic inhibition with a brain-penetrant IDO1 inhibitor restore hippocampal glucose metabolism and spatial memory not only in preclinical models of amyloid-ß accumulation but also in a model of tau accumulation, suggesting that these distinct pathologies share a common metabolic deficit.

## Results

### IDO1 inhibition restores astrocytic bioenergetic responses to amyloid Aß42 and Tau oligomers

We investigated whether IDO1 activity functioned in astrocytic responses to amyloid-ß_42_ oligomers (oAß) and Tau oligomers (oTau), the two major neurotoxic pathologies in AD. Mouse primary astrocytes showed prominent IDO1-dependent KYN production in response to oAß, oTau, and the combination of oAß+oTau (**Supplementary Fig. 1A-C**, **Fig. 1A-B**). We also observed AD pathology-induced activation of IDO1 and KYN production in human iPSC-derived astrocytes (iAstrocytes) (*14*) (**Fig. 1C, Supplementary Fig. 1D**), demonstrating a conserved IDO1 response to Aß and Tau oligomers across mouse and human astrocytes. IDO1 generation of KYN was blocked in all cases with the selective IDO1 inhibitor PF06840003 (PF068) (*15*). To confirm that oAß+oTau increased KYN specifically via activation of IDO1, we administered PF068 in combination with ^13^C-labeled-tryptophan (TRP) and traced KYN levels in mouse and human astrocytes (**Figure 1D; Supplementary Fig. 1E-G**). Consistent with activation of astrocytic IDO1, high levels of TRP-derived KYN were generated by astrocytic IDO1 in response to oAß+oTau in both mouse and human astrocytes.

**Figure 1.**
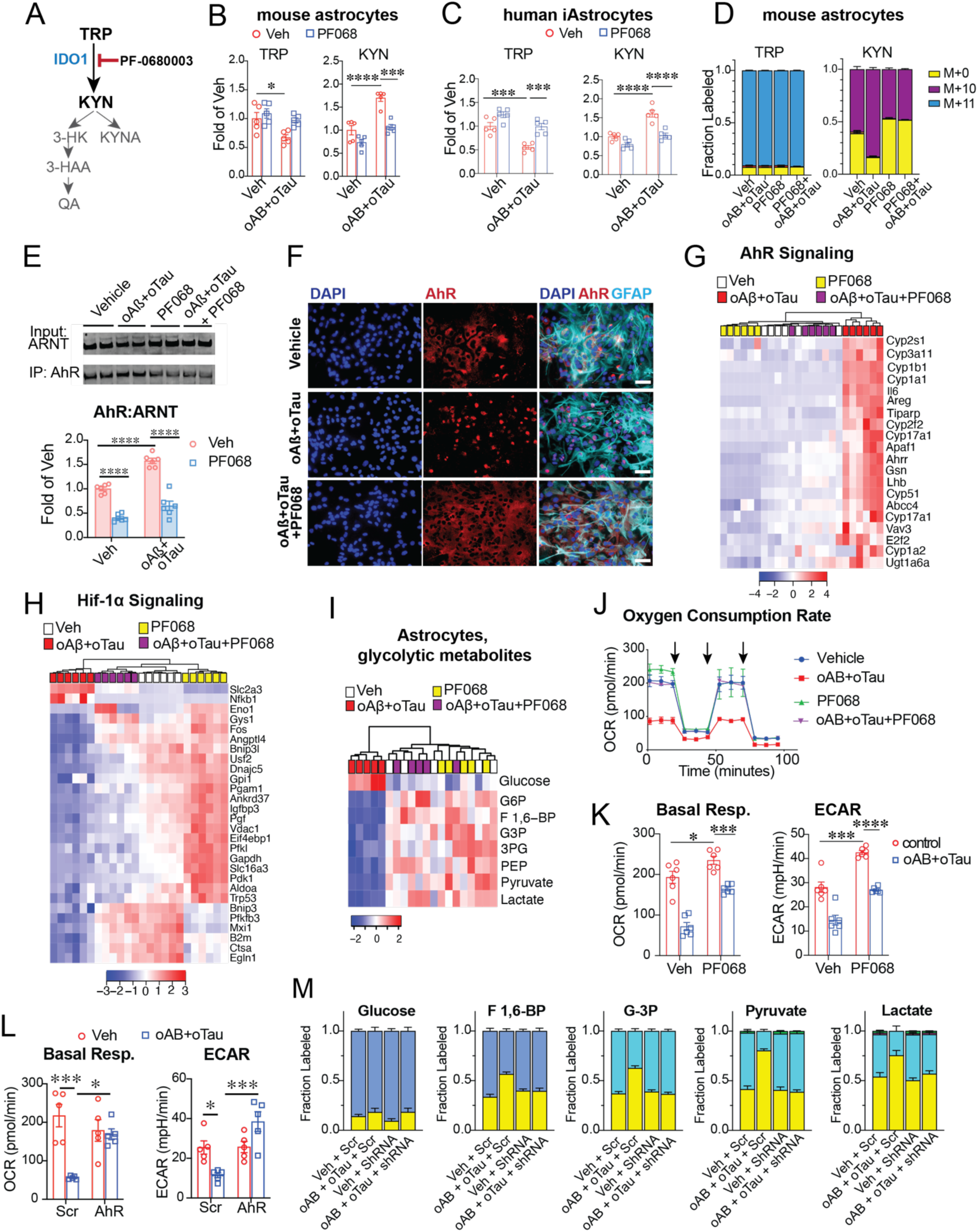
Activation of astrocytic IDO1 in response to amyloid Aß_42_ and/or tau oligomers suppresses astrocytic glycolysis. Data are mean ± s.e.m. and analyzed using two-way ANOVA with Tukey post hoc tests: \**P*<0.05, \*\**P*<0.01, \*\*\**P*<0.001 and \*\*\*\**P*<0.0001, unless otherwise specified. **(A)** The kynurenine pathway (KP): the essential amino acid tryptophan (TRP) is metabolized to kynurenine (KYN) by IDO1. The specific IDO1 inhibitor PF0680003 (PF068) prevents KYN production. **(B)** Primary mouse astrocytes were stimulated with veh or oAβ+oTau (100nM, 20h) +/- PF068 (100nM, 20h). LC-MS quantification of TRP and KYN (n=5/group). **(C)** iPSC derived human iAstrocytes were stimulated with veh or oAβ+oTau (100 nM, 20h) +/- PF068 (100 nM, 20h). LC-MS quantification of TRP and KYN (n=5/group). **(D)** Isotope tracing of ^13^C-TRP [M+11] to KYN [M+10] in mouse astrocytes (n=5/group). Similar mass labeling occurs in human astrocytes (**Supplementary Fig. 1G**) **(E)** Coimmunoprecipitation (CoIP) of AhR and ARNT in mouse astrocytes. (Top) Representative immunoblot. (Bottom) Quantification of co-immunoprecipitated AhR:ARNT in astrocytes treated with oAβ+oTau +/- PF068 (100 nM, 20h). **(F)** Immunocytochemical detection of AhR (red) in primary mouse astrocytes (GFAP, blue) shows nuclear localization following stimulation with oAβ+oTau (100 nM, 20h) that is prevented with PF068 (100 nM, 20h). Scale bars=50µM. Quantification in **Supplementary Fig. 2B**. **(G)** qRT-PCR of AhR-dependent gene transcripts significantly altered in mouse astrocytes +/- oAβ+oTau (*q* < 0.05). **(H)** qRT-PCR of Hif-1α-dependent gene transcripts significantly altered in mouse astrocytes +/- oAβ+oTau (*q* < 0.05). **(I)** LC/MS of glycolytic metabolites in primary astrocytes +/- oAβ+oTau (100 nM, 20h) and +/- PF068 (100 nM, 20h); n=5/group). **(J)** Realtime oxygen consumption rate (OCR) in mouse astrocytes +/- oAβ+oTau (100 nM, 20h) +/- PF068 (100nM, 20h; n=6/group). Cells were treated with 1 μM oligomycin (olig), 2 μM carbonyl cyanide-p-trifluoromethoxyphenylhydrazone (FCCP) and 0.5 μM rotenone and antimycin (rot/an), as indicated by the three arrows. **(K)** Quantification of basal respiration and extracellular acidification rate (ECAR) from **(J)**, n=6/group. **(L)** Mouse astrocytes were transfected with either scrambled shRNA (Scr) or shRNA to AhR (AhR). Quantification of basal respiration (left) and ECAR (right), n=5/group. **(M)** Isotope-tracing of ^13^C-glucose administration in mouse astrocytes transfected with either shRNA (Scr) or shRNA to AhR and stimulated with veh or oAβ+oTau (100 nM, 20h; n=5/group). F 1,6-BP = Fructose 1,6-bisophosphate, G-3-P = glyceraldehyde 3-phosphate.

We then investigated the downstream effects of IDO1 activation in astrocytes. KYN binds to the AhR and triggers translocation of the AhR to the nucleus (*11, 16*). In the nucleus, the AhR dimerizes with the AhR nuclear translocator (ARNT) to regulate transcriptional responses (**Supplementary Fig. 2A**). We tested whether stimulation of astrocytes with oAß+oTau induced IDO1-dependent AhR:ARNT binding. Co-immunoprecipitation of AhR and ARNT increased in response to Aß_o_+Tau_o_ and was inhibited by the IDO1 inhibitor PF068 (**Fig. 1E**), indicating that oAß+oTau activates IDO1 and induces AhR:ARNT binding. Orthogonal validation using immunohistochemical detection of the AhR in astrocyte nuclei confirmed that IDO1 promotes AhR nuclear translocation (**Fig. 1F; Supplementary Fig. 2B**).

In the nucleus, ARNT levels are limiting because ARNT can also bind to HIF-1α (hypoxia-inducible factor 1-alpha; **Supplementary Fig. 2A**)(*17*), a transcription factor that regulates glycolytic gene transcription and increases generation of lactate (*18*). To assess astrocyte AhR/ARNT versus HIF1α/ARNT gene expression, we examined AhR versus HIF1α transcriptional responses to Aß_o_+Tau_o_ stimulation in mouse astrocytes +/- PF068. As expected, astrocytic AhR gene transcription increased dramatically following Aß_o_+Tau_o_ stimulation but was significantly reduced with IDO1 inhibition (**Fig. 1G**). Conversely, IDO1 inhibition increased HIF1α-regulated glycolytic gene expression in resting astrocytes and reversed suppression of HIF1α regulated transcripts in Aß_o_+Tau_o_ stimulated astrocytes (**Fig. 1H; Supplementary Fig. 2C-D**). Indeed, IDO1 inhibition restored glycolytic intermediates and lactate in Aß_o_+Tau_o_ stimulated astrocytes to control levels (**Fig. 1I; Supplementary Fig. 2E**) and restored glycolysis (measured as Extracellular Acidification Rate, or ECAR) and mitochondrial oxidative phosphorylation (measured as Basal Respiration) (**Fig. 1J-K; Supplementary Fig. 3A**). Thus, astrocytic glucose metabolism and bioenergetics are disrupted by Aß_o_+Tau_o_ in an IDO1-dependent fashion.

To further confirm the role of IDO1 in suppressing astrocytic glycolytic responses to Aß_o_+Tau_o_, we transfected astrocytes with an shRNA to AhR. Quantitative immunoblotting confirmed knockdown of AhR protein levels (**Supplementary Fig. 3B**) and Seahorse analysis revealed that AhR knockdown prevented Aß_o_+Tau_o_ driven deficits in astrocytic glucose metabolism (**Fig. 1L; Supplementary Fig. 3C**). To orthogonally verify that AhR-knockdown rescued glucose metabolism in the context of Aß and tau pathology, we performed ^13^C-glucose isotope tracing and observed a rescue of glucose incorporation in IDO1-inhibited astrocytes stimulated with oAβ+oTau (**Fig. 1M; Supplementary Fig. 3D**). Thus, KYN-dependent AhR signaling suppresses glucose metabolism in astrocytes exposed to Aß_o_+Tau_o_. These data suggest that IDO1 may negatively affect critical astrocytic support of neurons in AD.

### IDO1 inhibition rescues long term memory in amyloid models of AD pathology

Astrocytes are the major repository of glycogen in the brain, and glycogen can be rapidly catabolized to lactate and exported to fuel neuronal mitochondrial respiration and synaptic activity (*7–9*). Learning and memory are enabled by astrocyte-derived lactate (*9, 19–22*). Given the importance of astrocytic lactate in metabolic support of neurons, we hypothesized that IDO1 inhibition in astrocytes might improve hippocampal memory deficits and synaptic plasticity in models of AD pathology.

We first confirmed absence of IDO1 activity in primary neurons as compared to astrocytes. IDO1 inhibition with PF068 elicited a dose response increase in glycolysis and mitochondrial respiration in astrocytes but not neurons (**Fig. 2A-B**). We then assessed in vivo effects of PF068, which is brain penetrant, on hippocampal KYN and lactate levels in 6-month old wild type (WT) mice. IDO1 inhibition reduced hippocampal KYN and reciprocally increased lactate in a dose-dependent manner (**Fig. 2C**). We then tested the effects of PF068 in two mouse models that accumulate misfolded Aß_42_ peptides - the *APP^Swe^-PS1^ΔE9^* (APP/PS1) (*23*) and the 5X FAD (*24*) models. Hippocampal KYN levels were significantly increased in both mutant APP models and this effect was blocked with PF068, indicating that IDO1 activation occurs with Aß_42_ accumulation in vivo (**Fig. 2D-E; Supplementary Fig. 4C-D**). Hippocampal lactate was reduced in both models in an IDO1-dependent manner (**Fig. 2F-G**). These findings suggest a role for IDO1 and KYN generation in regulating hippocampal glucose metabolism in mouse models of amyloid accumulation.

**Figure 2.**
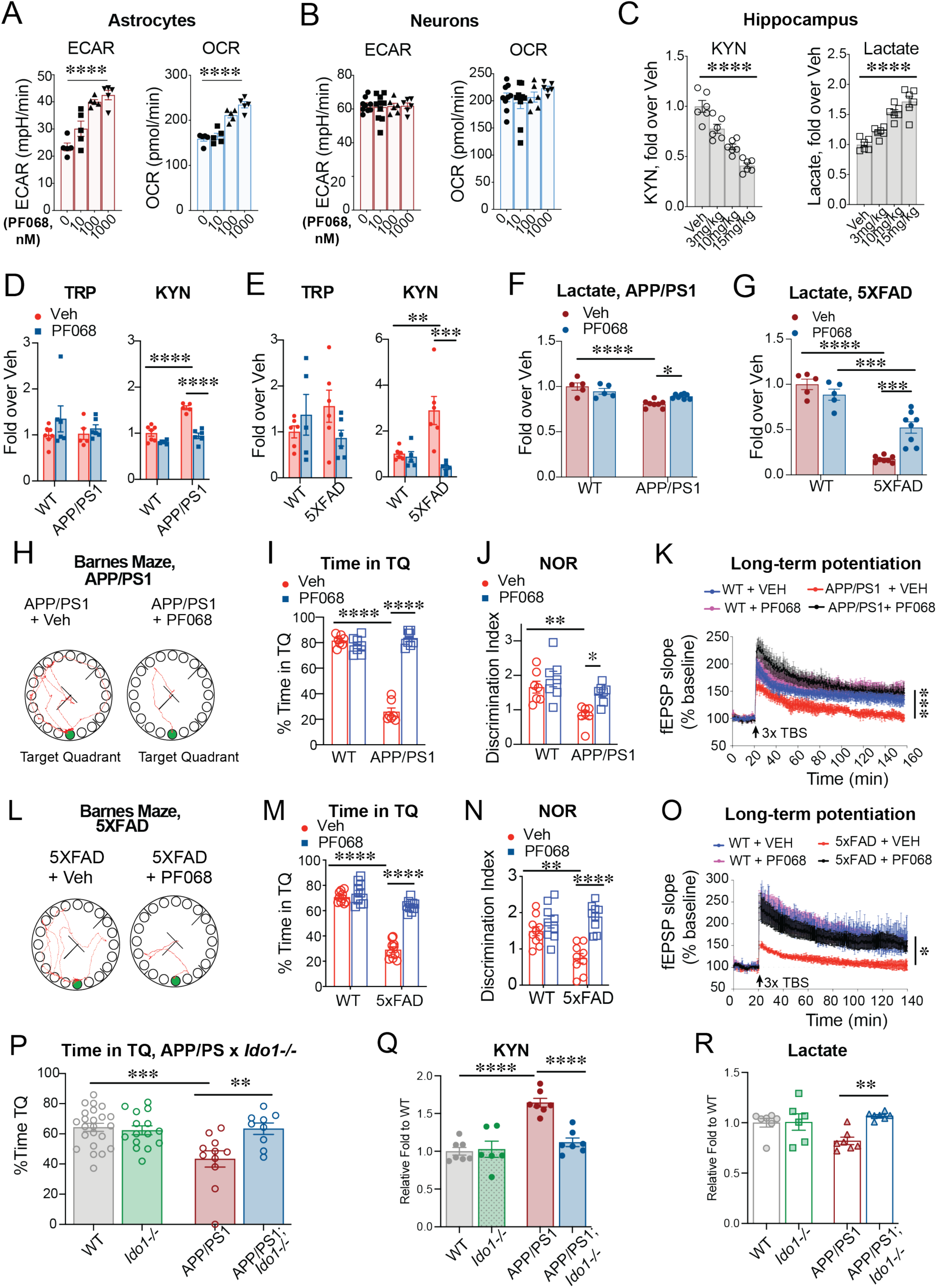
IDO1 inhibition restores hippocampal lactate levels, spatial memory, and long-term potentiation in mutant APP models. Data are mean ± s.e.m. and analyzed using two-way ANOVA with Tukey post hoc tests: \**P*<0.05, \*\**P*<0.01, \*\*\**P*<0.001 and \*\*\*\**P*<0.0001, unless otherwise specified. Wild type and APP/PS mice (10-12 mo males) and wild type and 5XFAD mice (5-6 mo females) were administered PF068 at 15 mg/kg/day for one month by gavage; mice were then behaviorally trained and tested or tested by electrophysiology. **(A)** Real-time changes in ECAR and OCR in mouse astrocytes stimulated with increasing doses of PF068 (n=5/group, 20h). One-way ANOVA, repeated measures, Tukey’s post-hoc test, \*\*\*\**P*<0.001. **(B)** Real-time changes in ECAR and OCR in mouse hippocampal neurons stimulated with increasing doses of PF068 (n=6-9/group, 20h). **(C)** LC-MS quantification of hippocampal KYN and lactate in 6 mo old WT mice (n=6 mice/group) treated with increasing doses of PF068 for 4h. One-way ANOVA, repeated measures, Tukey’s post-hoc test, \*\*\*\**P*<0.001. **(D)** Hippocampal TRP and KYN levels in APP/PS1 mice +/- PF068 at 15 mg/kg for one month (n=5-7/group). **(E)** Hippocampal TRP and KYN levels in 5xFAD mice +/- PF068 at 15 mg/kg for one month (n=5-6/group). **(F-G)** Hippocampal lactate levels in APP/PS1 mice (**F**) and 5X FAD mice (**G**) administered vehicle or 15mg/kg PF068 for 1 month. **(H)** Representative tracings on the day of testing in the Barnes maze of APP/PS1 mice +/- PF068 (n=7/group). The Target Quadrant (TQ) escape hole is in green. **(I)** Quantification of time in the target quadrant (TQ). **(J)** Discrimination index in the Novel Object Recognition (NOR) task for the number of interactions with the novel object in APP/PS1 mice +/- PF068 (n=7/group). **(K)** Long-term potentiation, measured as the change in field excitatory postsynaptic potential (fEPSP), in the CA1 hippocampal region over 160 min in APP/PS1 mice +/- 1 month of PF068 treatment. Three episodes of theta-burst stimulation (3 × TBS; black arrows) were applied. Two-way ANOVA, effects of time and genotype: *P* < 0.0001; Sidak’s multiple comparisons test with Geisser–Greenhouse correction \*\*\**P*<0.001 (n = 8-9 slices, 4-5 mice per group). **(L)** Representative tracings of the test trial day in the Barnes Maze in 5XFAD mice +/- PF068 (n=9-10/group). **(M)** Percentage of time in the TQ from (**L**) **(N)** Discrimination index in the NOR task in 5XFAD mice +/- PF068 (n=9-10/group). **(O)** Long-term potentiation in APP/PS1 mice +/- PF068. Two-way ANOVA, effects of time and genotype *P* < 0.0001; Sidak’s multiple comparisons test with Geisser–Greenhouse correction ***P<0.05 (n = 8-9 slices, 4-5 mice per group). **(P)** Morris water maze testing and time in the TQ for APP/PS1 with *Ido1* deletion (n=11-23/group). **(Q)** KYN levels in hippocampus of APP/PS1 +/- *Ido1* deletion (n=6-7 mice per group). **(R)** Lactate levels in hippocampus of APP/PS1 +/- *Ido1* deletion (n=6-7 mice/group).

We then tested whether IDO1 activity contributed to the well-established spatial memory and synaptic deficits observed in mutant APP models. IDO1 inhibition with PF068 reversed deficits in hippocampus-dependent spatial memory in the Barnes maze task and Novel Object Recognition in both the APP/PS1 and 5X FAD mouse models (**Fig. 2H-J and L-N; Supplementary Fig. 5A-C and E-G**) in the absence of significant changes in amyloid load (**Supplementary Fig. 5 J-K**). We then assessed hippocampal synaptic plasticity with electrophysiological recordings of the CA3 to CA1 Schaffer collateral pathway. In line with the behavioral findings, IDO1 inhibition also rescued deficits in long-term potentiation (LTP), a cellular correlate of learning and memory in both mutant APP models (**Fig. 2K and 2O; Supplementary Fig. 5D and 5H**). We further confirmed our pharmacologic findings genetically in APP/PS1;*Ido1^-/-^* mice. Consistent with our pharmacologic findings, genetic loss of IDO1 also prevented the hippocampal increase in KYN and decrease in lactate and rescued spatial memory function in the Morris Water Maze task (**Fig. 2 P-R**).

### IDO1 inhibition rescues metabolic and hippocampal memory changes in a tau model of AD pathology

Along with accumulation of amyloid, which is considered necessary but not sufficient for AD development (*25*), the second major pathology in AD is accumulation and spread of misfolded tau. Accordingly, we tested whether the metabolic and functional rescue seen with IDO1 inhibition in mutant APP models would also occur in the P301S tau model of AD pathology, which expresses a mutant form of human microtubule-associated protein tau (MAPT)(*26*). Indeed, hippocampal KYN was significantly increased and lactate was conversely decreased in this model as well (**Fig. 3A-B; Supplementary Fig. 6A**). Similar to our findings in mutant APP models, IDO1 inhibition in P301S tau mice rescued deficits in spatial memory in the Barnes Maze and NOR (**Fig. 3 C-D; Supplementary Fig. 6B-F**) and restored hippocampal LTP (**Fig. 3E; Supplementary Fig. 6G**). The striking rescue of hippocampal function across both amyloid and tau pathologies suggests a common underlying metabolic defect triggered by both pathologies and driven by IDO1-generated KYN.

**Figure 3.**
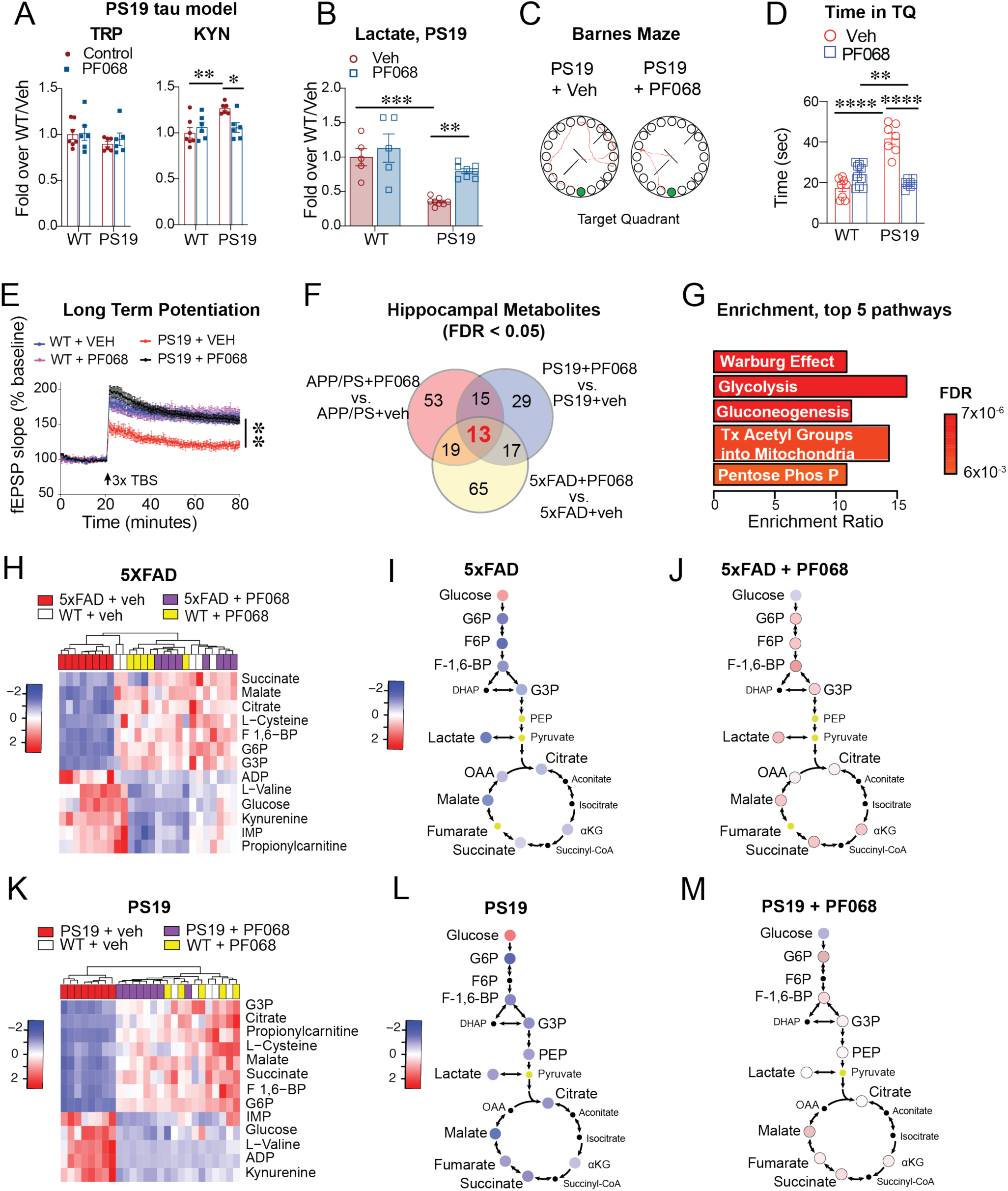
IDO1 inhibition rescues hippocampal function and glucose metabolism across amyloid and tau pathologies. PS19 and WT littermates (8-9 months old, n=7-10/group, males and females) were administered 15 mg/kg/day PF068 or vehicle for 1 month and then behaviorally or electrophysiologically tested. For metabolic assessments, APP/PS1 and WT littermates (10-12 months old), 5xFAD and WT littermates (5-6 months old), and PS19 and WT littermates were treated with veh or PF068 at 15 mg/kg/day for one month. Data are mean ± s.e.m. and are analyzed using two-way ANOVA with Tukey post hoc tests: \**P*<0.05, \*\**P*<0.01, \*\*\**P*<0.001 and \*\*\*\**P*<0.0001, unless otherwise specified. Hierarchical clustering is represented in terms of distance from the mean, or Z-score. **(A)** LC-MS quantification of TRP and KYN in hippocampus of PS19 mice (n=6/group) **(B)** Lactate levels in hippocampus of PS19 mice (n=5/group) **(C)** Representative tracings of path to escape hole (green) in PS19 mice +/- PF068 treatment on the day of testing in the Barnes maze. **(D)** Percentage of time in the TQ in PS19 mice. **(E)** Long-term potentiation in PS19 mice +/- PF068 treatment. Two-way ANOVA, effects of time and genotype *P* < 0.0001; Sidak’s multiple comparisons test with Geisser–Greenhouse correction **P<0.01 (n = 8-9 slices, 4-5 mice per group). **(F)** Venn diagram depicting number of significant metabolites (*q* < 0.05) detected by untargeted metabolomics from hippocampi in PF068-treated vs. veh-treated APP/PS1, 5XFAD, and PS19 mice. 13 metabolites are shared across the three comparisons of amyloid and tau pathologies. (**G**) Enrichment pathway analysis of 13 shared metabolites from (**F**) using MetaboAnalyst. (**H**) Hierarchical clustering of the 13 shared hippocampal metabolites in 5XFAD and WT littermates +/- PF068 (Z-score, -2 to 2). (**I-J**) Schematic depicting levels of glycolytic and TCA metabolites and their average Z-score value from **(H)**. Note the rescue of glycolysis and TCA with IDO1 inhibition in 5XFAD mice. (**K**) Hierarchical clustering of the 13 shared hippocampal metabolites in PS19 and WT littermates +/- PF068 (Z-score, -2 to 2). (**L-M**) Schematic depicting levels of glycolytic and TCA metabolites and their average z-score value from **(K)**. Note the rescue of glycolysis downstream of glucose with IDO1 inhibition in PS19 mice.

### IDO1 disrupts hippocampal glucose metabolism across amyloid and tau pathology models

To better understand the metabolic basis underlying the beneficial effects of IDO1 inhibition, we performed metabolomic analyses of hippocampus from the three preclinical models. We identified thirteen metabolites shared across the three models that demonstrated enrichment for glucose metabolism (**Fig. 3F-G)**. Remarkably, across both amyloid and tau pathologies, IDO1 inhibition restored multiple glycolytic and citric acid cycle (TCA) intermediates to WT levels, consistent with rescue of an energy-depleted state (**Figure 3H-M**; **Supplementary Fig 7A-C**; **Supplementary Fig. 8**). APP/PS1*;Ido1^-/-^* mice also demonstrated a significant rescue in hippocampal glucose metabolism as compared to APP/PS1 littermates (**Supplementary Fig. 7D**). Taken together, IDO1 inhibition - across both amyloid and tau pathologies – restores hippocampal glucose metabolism and lactate production, spatial memory, and synaptic plasticity.

To further confirm that IDO1 inhibition rescues active glucose metabolism in vivo, we carried out an orthogonal approach and infused isotope labeled ^13^C-glucose at a physiologic rate and concentration through the carotid artery of APP/PS1 and PS19 mice treated with vehicle or IDO1 inhibitor. We measured incorporation of labeled glucose into hippocampal glycolytic and TCA metabolites. IDO1 inhibition restored hippocampal glycolysis to WT levels across both amyloid and tau pathologies and restored glucose incorporation into downstream lactate and TCA intermediates (**Figure 4A-B**). To visually confirm these changes, we performed matrix-assisted laser desorption ionization (MALDI) imaging (*27, 28*). Imaging of vehicle and PF068-treated AD mice demonstrated significant declines in glucose incorporation into glycolytic (fructose-1,6 biphosphate and lactate) and TCA (malate) intermediates in both APP/PS1 amyloid and P301S tau models. These declines were reversed across both pathologies by IDO1 inhibition with PF068 (**Figure 4C-F**).

**Figure 4.**
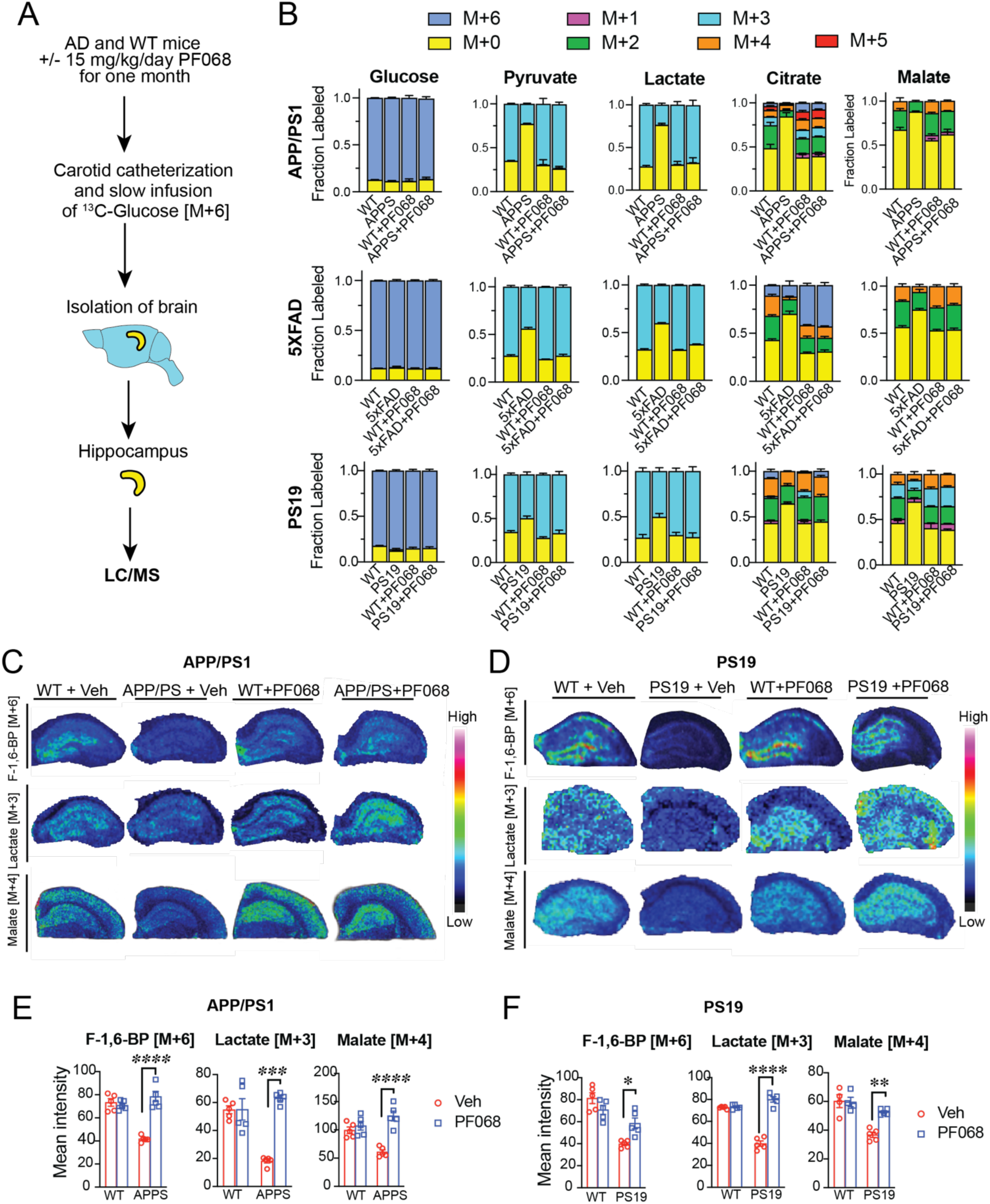
In vivo mass-labeling reveals that IDO1 inhibition rescues glucose incorporation into the TCA cycle in hippocampi of AD model mice. APP/PS1 mice (10-12 months old, n=5/group), 5xFAD mice (5-6 months old, n=5/group) and PS19 mice (8-9 months old, n=5/group) and age-matched WT littermates were treated +/- PF068 for one month. Mice subsequently underwent heavy isotope-labeling experiments and MALDI imaging. Data are mean ± s.e.m. and analyzed using two-way ANOVA with Tukey post hoc tests: \**P*<0.05, \*\**P*<0.01, \*\*\**P*<0.001 and \*\*\*\**P*<0.0001, unless otherwise specified. (**A**) Schematic depicting carotid catheterization and infusion of ^13^C-glucose to achieve steady-state isotope labeling of glucose *in vivo* in AD model mice. (**B**) Isotope tracing of ^13^C-glucose metabolism in hippocampi of APP/PS1 (top row), 5XFAD (middle row), and PS19 (bottom row) mice treated with veh or PF068. IDO1 inhibition with PF068 restores glucose incorporation into glycolytic and TCA metabolites across both amyloid and tau pathologies. (**C-D**) Representative images of matrix-assisted laser desorption/ionization (MALDI) of coronal hippocampal sections from APP/PS +/-PF068 and PS19 +/-PF068 mice showing rescue in ^13^C-labeled glycolytic intermediate fructose-1,6-BP [M+6], lactate [M+3], and TCA intermediate malate [M+4]. (**E-F**) Mean fluorescent intensity (MFI) quantification of **C-D**.

### IDO1 inhibition rescues lactate-dependent LTP

Given our in vitro data demonstrating a role for IDO1 in regulating astrocytic lactate production, we hypothesized that restoration of hippocampal LTP in PF068-treated AD mice might be dependent on lactate produced by astrocytes that is then transferred to neurons. Accordingly, we administered an inhibitor of monocarboxylate transporters (MCT) MCT1 and MCT2 (AR-C155858) (*29*) to hippocampal slices derived from AD mice treated with vehicle or IDO1 inhibitor (**Fig. 5A-C**). Inhibition of MCT1/2 will disrupt lactate translocation from the astrocyte to neuron (**Fig 5D**). MCT1/2 inhibition prevented the PF-068-mediated rescue of LTP in all three AD models, suggesting that IDO1-inhibition rescues LTP in a lactate-dependent manner across distinct pathologies.

**Figure 5.**
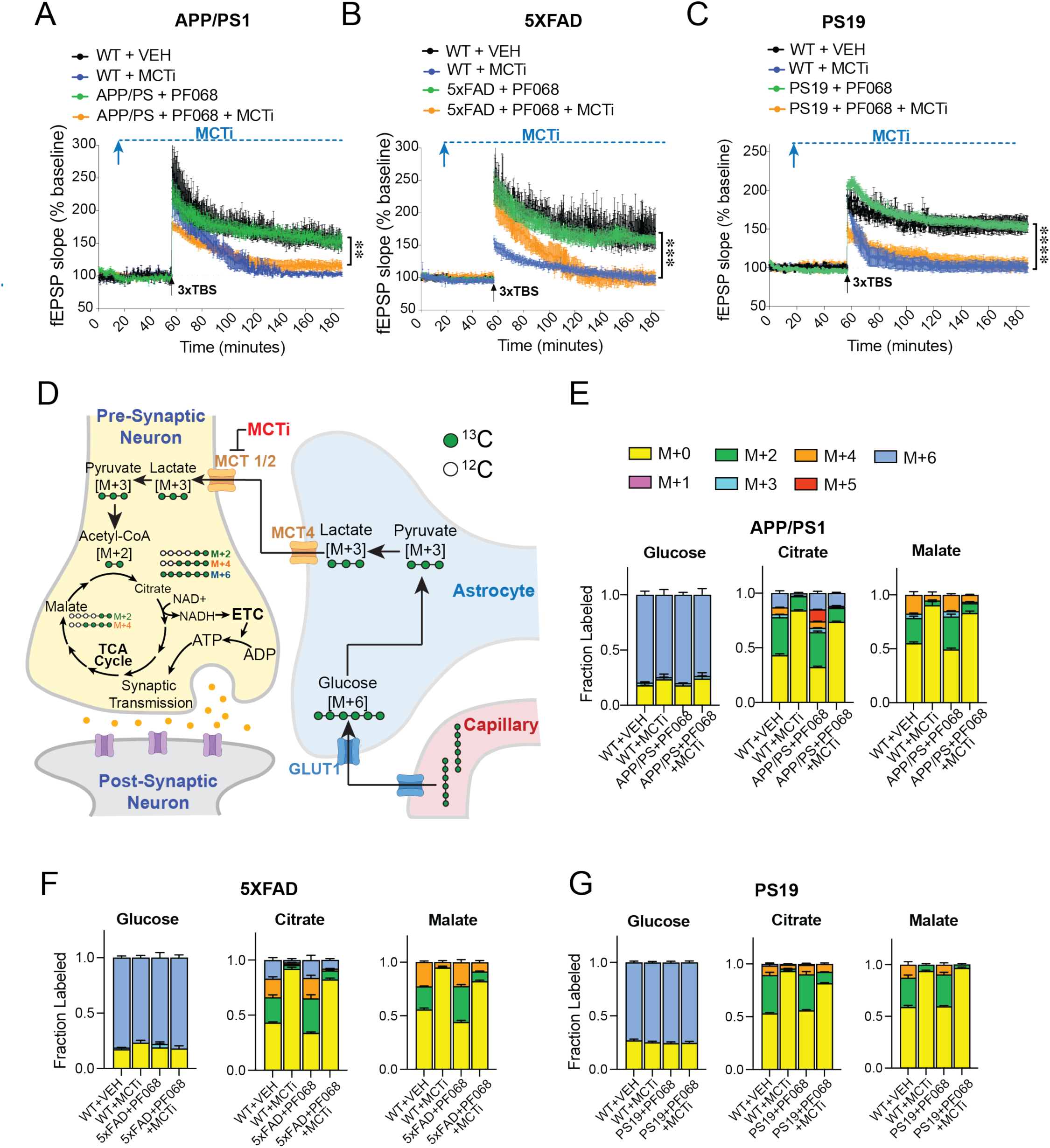
IDO1 inhibition rescues lactate-dependent LTP in hippocampi of AD model mice. APP/PS1 mice (10-12 months old), 5xFAD mice (5-6 months old) and PS19 mice (8-9 months old) and WT littermates were treated +/- PF068 at 15 mg/kg/day for one month and then hippocampal slices were assessed by electrophysiology and mass labeling. Data are mean ± s.e.m. (**A-C**) Long-term potentiation, measured as the change in field excitatory postsynaptic potential (fEPSP) in the CA1 hippocampal region over 180 min. Hippocampal slices from PF068-treated AD and WT mice were stimulated with monocarboxylate transporter-1/2 inhibitor (MCTi, 50µM) beginning 15 minutes after the start of the recording (blue arrow) and 45 minutes before theta-burst stimulation (3 × TBS; black arrows) and continuing for the whole recording interval. Two-way ANOVA, effects of time and genotype *P* < 0.0001; Sidak’s multiple comparisons test with Geisser–Greenhouse correction \*\**P*<0.01, \*\*\**P*<0.001, \*\*\*\**P*<0.0001 (n = 8-9 slices/hippocampus). **(A)** Comparison between APP/PS+ PF068 and APP/PS+PF068+MCTi, \*\**P*<0.01 (n=5 mice/group). **(B)** Comparison between 5XFAD+ PF068 and 5XFAD+PF068+MCTi, \*\*\**P*<0.001 (n=6 mice/group). **(C)** Comparison between PS19+ PF068 and PS19+PF068+MCTi, \*\*\*\**P*<0.0001 (n=6-8 mice/group). (**D**) Schematic detailing tracing of ^13^C-glucose labeling from the capillary to the neuron. The capillary contains [M+6] glucose. [M+6] glucose is taken up by the astrocyte foot process and metabolized to [M+3] lactate, which is then transported out of the astrocyte by MCT4 and into the neuron by MCT1/2. In the neuron, [M+3] lactate is converted to [M+3] pyruvate and then enters the TCA to fuel oxidative phosphorylation for generation of ATP for synaptic transmission. (**E-G**) Isotope tracing of ^13^C-glucose in hippocampal slices that had undergone LTP derived from APP/PS1 (**E**) 5XFAD (**F**) and PS19 (**G**) mice. A rescue in glucose incorporation into the TCA cycle occurs in hippocampi from AD model mice treated with IDO1 inhibitor. This rescue is blocked with administration of MCT1/2 inhibitor (n=5 mice/group, MCTi, 50µM).

We also performed ^13^C-glucose mass labeling of hippocampal slices from AD model mice treated with PF068 that had undergone electrophysiology testing. Here, we examined downstream incorporation of labelled glucose into TCA intermediates in the presence or absence of MCT1/2 inhibitor (**Fig. 5D-G**). As shown previously with in vivo mass labeling in **Figure 4**, IDO1 inhibition in APP/PS1, 5X FAD, and PS19 hippocampal slices showed restoration of mass-labelled citrate and malate to WT levels, consistent with restoration of glucose flux into the TCA. However, this rescue was completely prevented with MCT1/2 inhibition, suggesting that synaptically active neurons utilize lactate in an IDO1-dependent manner across both amyloid and tau pathologies. Thus, inhibition of IDO1 in amyloid and tau models rescues lactate production and uptake into neurons which, in turn, can rescue ATP-dependent synaptic activity.

### IDO1 inhibition restores lactate transfer from hAstrocytes to hNeurons from subjects with late-onset AD (LOAD)

The APP/PS1, 5X FAD, and PS19 models recapitulate the individual pathologies of rare genetic forms of familial AD (FAD) and tauopathy. The majority of AD, however, is late-onset (LOAD), with more complex etiologies. To investigate whether KYN levels are altered in LOAD, we measured KYN and TRP levels in well-characterized human post-mortem brain tissues that harbored increasing levels of Braak pathology, the basis for neuropathological diagnosis of AD (*30*). KYN and TRP levels were assessed in the middle frontal gyrus, a brain region that demonstrates bilateral gray matter loss (*31*), synaptic loss, and high Aβ burden (*32*). The tissue was obtained from subjects classified as: non-demented (Braak I–II, n = 12), demented Braak I–II (non-AD, n = 12), AD Braak III–IV (AD-mid, n = 12) and AD Braak V–VI (AD-high, n = 12). Diagnostic groups were balanced for age, gender, and post-mortem interval. Demographic information of the cohort is shown in **Supplementary Table 1**. LC/MS revealed a significant increase in KYN but not TRP with increasing Braak stage (**Fig. 6A**).

**Figure 6.**
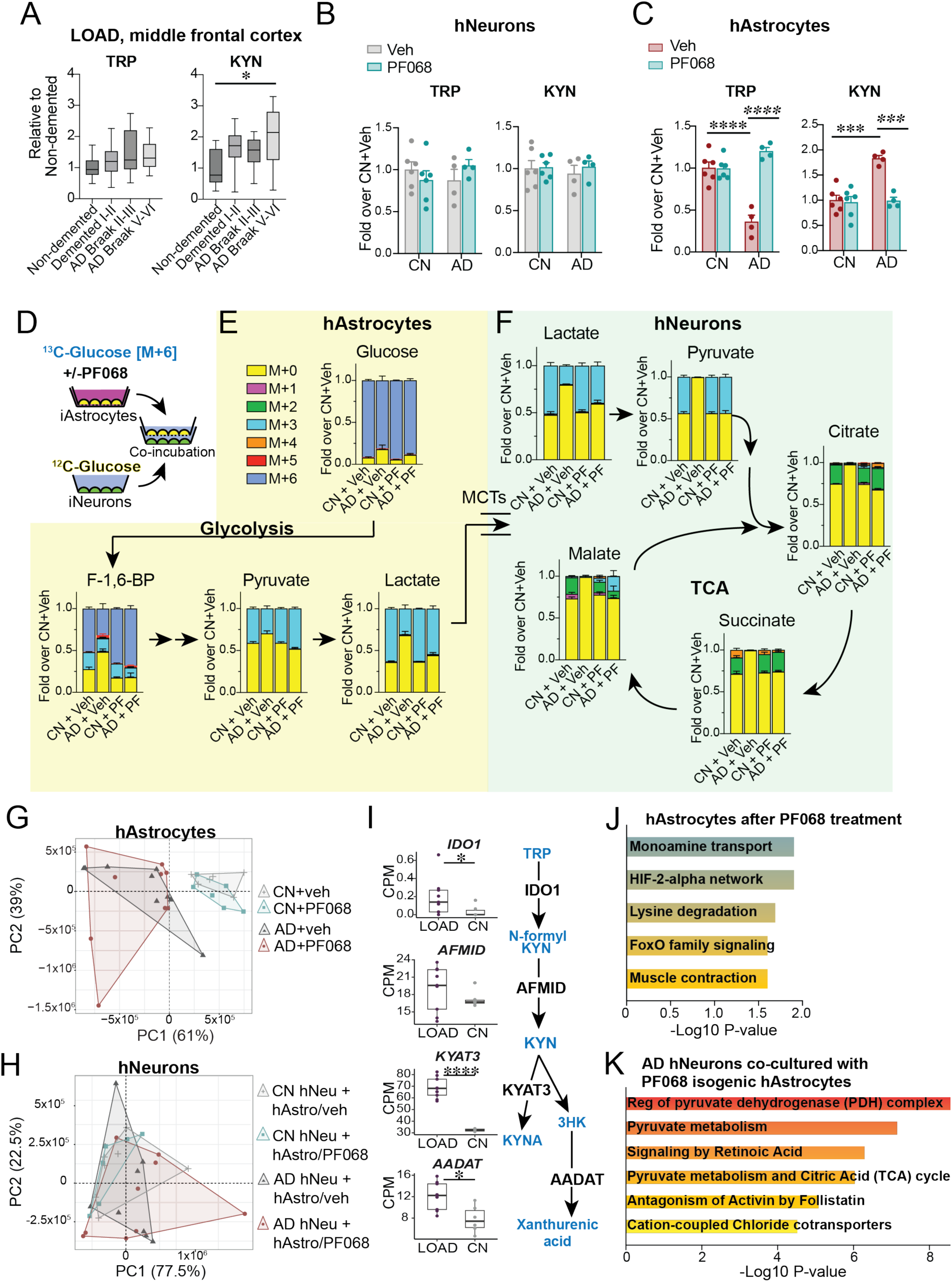
IDO1 inhibition restores lactate transfer from hAstrocytes to hNeurons in late-onset AD. **(A)** LC-MS quantification of TRP and KYN in middle frontal gyrus from non-demented Braak I-II, demented non-AD Braak stages I-II, AD Braak III-IV and AD Braak V-VI brain (n=12 donors per group). One way ANOVA, \**P*<0.05. **(B)** LC-MS quantification of TRP and KYN in cognitively normal (CN) and AD hNeurons (n=6 CN, n=4 AD). **(C)** LC-MS quantification of TRP and KYN in CN and AD hNeurons (n=6 CN, n=4 AD). 2-way ANOVA with Tukey’s post-hoc test, \*\*\**P*<0.001, \*\*\*\**P*<0.0001. **(D)** Schematic depicting experimental design for mass labeling of hAstrocytes and subsequent co-culture with hNeurons. hAstrocytes were paired with isogenic hNeurons derived from n=4 AD and n=6 CN subjects. hAstrocytes from CN and AD patients were labeled with ^13^C-glucose and stimulated with veh or PF068 (100nM, 20h). hAstrocytes on well-inlets were then washed, transferred and co-incubated with neurons for 4h. LC/MS was then performed on cell lysates of hAstrocytes and hNeurons. **(E)** Isotope tracing of ^13^C-glucose in hAstrocytes +/- PF068 (100nM, 20h) shows rescue of glucose incorporation into glycolytic intermediates and [M+3] lactate (n=6 CN, n=4 AD) **(F)** ^13^C-labelled hAstrocytes from (**E**) were washed, transferred, and co-incubated with hNeurons for 4 hours. Mass-labelled astrocytic [M+3] lactate is taken up by hNeurons via the MCT1/2 transporter, and [M+3] lactate metabolism and incorporation into hNeuron pyruvate and TCA intermediates was measured. Glucose flux is restored to CN levels in AD hNeurons co-cultured with hAstrocytes that had been treated with PF068 (n=6 CN, n=4 AD). **(G)** Principal component analysis of RNA-seq DEGs from hAstrocytes derived from CN and AD subjects +/- PF068 (100 nM, 20h; n=2 CN and n=3 AD, technical triplicates). **(H)** Principal component analysis of DEGs from CN and AD hNeurons that were co-cultured with congenic hAstrocytes stimulated +/- PF068 (100 nM, 20h; n=2 CN and n=3 AD, technical triplicates) **(I)** (Left) Box plots of kynurenine pathway (KP) transcripts from hAstrocytes derived from CN or LOAD subjects. Wald test, * *P*<0.05, \*\*\*\**P*<0.001 (n=2 CN and n=3 AD, technical triplicates). (Right) Diagram of KP highlighting enzymatic steps corresponding to transcripts increased in LOAD hAstrocytes. **(J)** Top significantly enriched gene ontology terms in LOAD hAstrocytes of transcripts significantly regulated by PF068 versus vehicle. **(K)** Top significantly enriched gene ontology terms from differentially regulated transcripts in LOAD hNeurons co-cultured with isogenic hAstrocytes that had been treated with PF068 versus vehicle. Note enrichment of pyruvate and TCA metabolic pathways.

Next, we asked if the effect of IDO1 on the regulation of glucose metabolism could be modeled in LOAD human astrocytes and neurons in vitro. Patient specific human iPSC lines were differentiated into pairs of genetically matched neurons (hNeurons) and astrocytes (hAstrocytes) (**Supplementary Fig. 9A-B**). We performed LC-MS quantification of TRP and KYN from hNeurons and hAstrocytes derived from cognitively normal (CN) and LOAD subjects. We observed no changes in hNeuron TRP, KYN, or downstream KP metabolites between CN and LOAD subjects. However, hAstrocytes showed significant differences, with TRP significantly reduced and KYN significantly increased in LOAD hAstrocytes as compared to CN hAstrocytes (**Fig. 6B & C; Supplementary Fig. 9C-D**). Application of the IDO1 inhibitor PF068 to LOAD astrocytes restored TRP and KYN to CN levels, indicating increased IDO1 activity in LOAD astrocytes as compared to CN astrocytes.

We then tested the effect of IDO1 inhibition on the regulation of glucose metabolism and lactate transfer in hAstrocytes co-incubated with hNeurons from CN and LOAD subjects. hAstrocytes were mass labelled with ^13^C-glucose and treated with either vehicle or PF068 for 20 hours. The hAstrocytes were then washed, and transferred into a co-culture with their respective isogenic hNeurons for four hours to allow the hAstrocyte-generated [M+3] lactate to be taken up by hNeurons (**Fig. 6D**). In hAstrocytes from LOAD subjects, glucose metabolism was significantly reduced as compared to CN hAstrocytes, however this deficit was reversed to CN levels following IDO1 inhibition (**Fig 6E; Supplementary Fig. 9E**). In particular, [M+3] lactate levels in AD hAstrocytes, which were significantly reduced compared to CN, were normalized to CN levels with PF068. Upon co-incubation of PF068-treated hAstrocytes with hNeurons, [M+3] lactate uptake in hNeurons was restored to CN levels (**Fig. 6F; Supplementary Fig. 9F**). Co-incubation of LOAD hNeurons with PF068-treated hAstrocytes restored glucose incorporation into pyruvate and downstream TCA metabolites. Taken together, these findings suggest that IDO1 generation of KYN suppresses lactate production in LOAD hAstrocytes and subsequent uptake by hNeurons. IDO1 inhibition restored glucose flux and lactate uptake in LOAD hNeurons, replenishing the TCA that fuels mitochondrial respiration and synaptic activity.

The metabolic rescue induced by IDO1 inhibition in LOAD astrocytes and neurons resembled that observed in hippocampus of our preclinical mutant APP and Tau models. To identify potential mechanisms underlying this in vitro effect, we performed RNA sequencing on a subset of our LOAD and CN human iPSC cohort. Principle component analysis (PCA) indicated a robust disease status signature and mild drug effect in LOAD hAstrocytes, but no significant differences in hNeurons regardless of disease status or drug treatment (**Fig. 6G-H**). Differential gene expression was evaluated for every condition and revealed differentially expressed genes (DEGs) in vehicle-treated LOAD versus CN hAstrocytes that included genes encoding KP enzymes (**Fig. 6I**). While the transcriptional effect of PF068 was small, gene ontology analysis of LOAD neurons co-cultured with genetically matched, PF068-treated hAstrocytes demonstrated changes in pathways associated with increased pyruvate utilization and TCA function, similar to what we observed in preclinical models of AD (**Fig. 6J-K; Supplementary Fig, 10A-B**). Motif analysis on DEGs from comparisons between LOAD and CN hAstrocytes revealed a significant enrichment of the AhR-ARNT motif (**Supplementary Fig. 10C**), a signature that was not present in any other comparison. Unbiased hierarchical clustering of AhR-ARNT-associated transcripts identified by motif analysis demonstrated separation of AhR target genes between LOAD and CN hAstrocytes (**Supplementary Fig. 10D**), suggesting a role for the IDO1 pathway and KYN generation in regulating astrocytic responses in LOAD.

## Discussion

In this study, we combined in vitro and in vivo preclinical models to explore the role of astrocytes in AD pathogenesis. We identified the KP, and specifically IDO1-generated KYN, as a suppressor of astrocytic glucose metabolism and metabolic support of neurons. In vitro, Aß_42_ and tau oligomers disrupted astrocytic bioenergetics in an IDO1-dependent manner, where induction of KYN led to nuclear translocation of AhR and activation of AhR:ARNT gene transcription. Blockade of KYN production with an IDO1 inhibitor conversely activated HIF1α:ARNT transcription of glycolytic genes and restored astrocyte glucose metabolism and production of lactate. In vivo, mutant APP and tau transgenic models administered an IDO1 brain-penetrant inhibitor or APP/PS1 mice with a genetic deletion of *Ido1* showed reduced hippocampal KYN and reciprocally increased lactate. IDO1 blockade also reversed amyloid- and tau-mediated disruption of hippocampal glycolysis, spatial memory, and LTP. Confirmatory in vivo mass labeling of hippocampus with ^13^C glucose demonstrated that IDO1 inhibition and reduction of KYN restored glucose incorporation into lactate and TCA metabolites. Indeed, slice electrophysiology combined with mass labeling indicated that the rescue of LTP by IDO1 inhibition in APP/PS, 5XFAD, and P301S hippocampal slices was dependent on functional MCT1/2 transporters that carry lactate from astrocytes to neurons. IDO1-dependent suppression of astrocyte lactate production was also observed in human astrocytes derived from AD but not CN subjects. Mass labeling demonstrated that IDO1 inhibition of hAstrocytes restored the uptake and metabolism of astrocytic [M+3] lactate in co-cultured hNeurons. In human post-mortem frontal cortex, cerebral KYN levels rose with increasing Braak AD stages, in line with the observation of increased IDO1-derived hippocampal KYN in APP/PS, 5XFAD, and P301S preclinical models. Taken together, our data suggest that increased IDO1 activity and KYN generation across amyloid and tau pathologies suppress hippocampal metabolism, spatial memory, and synaptic plasticity. Restoring astrocytic glucose metabolism with IDO1 inhibition may represent an alternative therapeutic modality for neurodegenerative diseases like AD that is distinct from current amyloid lowering strategies.

The characteristic spatial distribution of glucose hypometabolism in parietal and temporal cortex by FDG-PET has enabled clinical differentiation of AD from other dementias. At a molecular level, recent proteomic studies provide some insight into this clinical observation by identifying disruptions in neuroglial glucose metabolism, notably in microglia and astrocytes (*5*). One limitation of our study is that we did not investigate the effects of IDO1 in brain microglia, where its expression is elevated in AD (*33*). KYN produced by microglia could potentially be taken up by astrocytes, where it would suppress astrocyte glucose metabolism. Alternatively, elevated KYN production in microglia could lead to increased generation of the downstream metabolite QA, an agonist of the N-methyl-D-aspartate (NMDA) receptor and a potential neurotoxin. Although one outcome of inhibiting IDO1 activity could be decreased generation of downstream neuroactive KP metabolites KYNA and QA (*34*), we did not observe PF068-dependent changes in these metabolites in vitro in astrocytes or neurons or in hippocampi from mutant APP and tau models. We observed a conserved effect of IDO1 inhibition on astrocytic glucose metabolism across the two major AD pathologies, amyloid and tau, in both in vitro and in vivo models.

Astrocytes perform essential functions, including facilitating synapse formation and plasticity, transporting glucose and other molecules from blood via the endothelium, immune surveillance, and maintenance of neurotransmitter pools. Astrocytes also provide fuel for neuronal activity in the form of lactate derived from glucose or glycogen (*20, 22, 35*). IDO1 blockade rescued synaptic potentiation across amyloid and tau pathologies in an MCT1/2-dependent fashion, indicating that astrocytic lactate transfer to neurons is regulated by astrocytic IDO1. We explored whether this relationship also occurred in congenic human astrocyte/neuron pairs derived from control and AD subjects. Here, astrocytes from AD subjects diverged away from control subjects in their production of KYN and in regulation of induction of KP and AhR-regulated transcripts. While we cannot explain why the AD signature was only observed in hAstrocytes and not hNeurons, we have controlled for purity of glial progenitor cells and hAstrocytes, and the yield and time to generate colonies were similar across CN and AD iPSC lines. As these experiments did not involve Aß42 or tau oligomers, the difference between AD and CN hAstrocytes may be cell intrinsic, potentially reflecting genetic or epigenetic differences between CN and AD disease states. Importantly, labeling with ^13^C-Glucose confirmed that IDO1 inhibition rescued AD hAstrocyte glycolysis and lactate generation. In co-cultures of human astrocytes and neurons derived from AD subjects, deficient astrocyte lactate transfer to neurons was corrected by IDO1 inhibition, resulting in improved neuronal glucose metabolism.

In addition to uncovering a novel role of IDO1 in brain glucose metabolism, our study highlights the potential of IDO1 inhibitors, developed as an adjunctive therapy for cancer, to be repurposed for neurodegenerative diseases like AD. This study also reveals a more general mechanism contributing to neuronal dysfunction that cuts across distinct pathologies. In addition to AD, manipulation of IDO1 may be relevant to Parkinson’s disease dementia (*36*), which is characterized by amyloid and tau deposits in addition to α-synuclein, as well as the broad spectrum of tauopathies. There is the possibility that deficient astrocytic glucose metabolism could also underlie other neurodegenerative diseases characterized by accumulation of additional misfolded proteins where increases in KP metabolites have been observed (*37*).

## Acknowledgements

This work was supported by RO1AG048232 (KIA), RF1AG058047 (KIA), P30 AG0066515 (KIA), American Heart Foundation (KIA 19PABH1345800 and FHG 19PABHI34610000), The Phil & Penny Knight Initiative for Brain Resilience at the Wu Tsai Neurosciences Institute, Stanford University (KIA), 5T32GM007365-45 (PSM), The Paul and Daisy Soros Fellowship for New Americans (PSM), the Gerald J. Lieberman Fellowship (PSM), HHMI Hanna H. Gray Fellows Program GT15655 (MRM), Burroughs Wellcome Fund PDEP 1022694 (MRM), the Scully Initiative (ALH, FML), AMED-Moonshot Program (Y.S and M.S), Wu Tsai Neurosciences Knight Initiative for Brain Resilience Scholar Award (TU). Data and tissue were obtained from the Arizona Study of Aging and Neurodegenerative Disorders (AZSAND), supported by NINDS U24 NS072026, NIA P30 AG19610, the Arizona Department of Health Services, the Arizona Biomedical Research Commission and the Michael J. Fox Foundation for Parkinson’s Research. We are grateful to the Stanford Behavioral and Functional Laboratory, the Stanford Mass Spectroscopy Core, and the Stanford Neuroscience Microscopy Service. Human iPSCs were derived by the Salk Stem Core Facility, supported by NIA P30 AG062429. We are grateful to the study participants and staff of the Shiley-Marcos UCSD Alzheimer’s Disease Center. KIA is a Chan Zuckerberg-San Francisco Biohub Investigator.

## SUPPLEMENTARY MATERIALS

I. **Methods and References**
II. **Supplementary Table 1**
III. **Supplementary Figures and legends**

## METHODS

### Animals

This study was conducted in accordance with National Institutes of Health (NIH) guidelines and the Institutional Animal Care and Use Committee at Stanford University approved protocols. All mice were housed in an environmentally controlled, pathogen-free barrier facility in an environment controlled for lighting (12-h light–dark cycle), temperature, and humidity, with food and water available ad libitum. The Stanford Veterinary Service Center monitors and maintains pathogen-free mouse housing as described in http://med.stanford.edu/vsc/about/rodenthandbook/rodent-diseases.html. C57B6/J congenic *APP^Swe^;PS1^ΔE9^*mice (*1*) were obtained from Jackson Labs (MMRRC Stock 34832) and overexpress a chimeric mouse/human amyloid precursor protein (Mo/HuAPP695swe) and a mutant human presenilin 1 (PS1-dE9). 5XFAD mice (*2*) were obtained from Jackson Labs (MMRRC Stock 34848) and overexpress mutant human amyloid beta (A4) precursor protein 695 (APP) with the Swedish (K670N, M671L), Florida (I716V), and London (V717I) Familial Alzheimer’s Disease (FAD) mutations and human PS1 with two FAD mutations, M146L and L286V. PS19 mice (*3*) were obtained from Jackson Labs (JAX stock #008169) and express the P301S mutant form of human microtubule-associated protein tau (*MAPT*). C57BL/6J *Ido1*-/- mice were obtained from Jackson laboratories (Stock No. 005867, Ido1^tm1Alm^, original contributor: Andrew Mellor, Medical College Georgia) and crossed with *APP^Swe^;PS1^ΔE9^*mice. Mice were dosed with PF06840003 or vehicle (PF068; Chemietek, CT-PF0684; stock solution 200 mM in DMSO) (*4*) at 15 mg/kg/day by gavage for 1 month.

### Neuron and astrocyte culture

For neuronal cultures, hippocampi were dissected from embryonic day C57BL/6 17.5 mouse embryos, dissociated using trypsin (2 mg/ml) and DNase I (0.6 mg/ml), and plated at a density of 100,000 cells per well in a Seahorse XF24 culture plate coated with poly-L-lysine. Neurons were maintained in Neurobasal medium, B27 (Invitrogen) and penicillin–streptomycin (Invitrogen) at 37 °C in a humidified atmosphere containing 5% CO2. The medium was refreshed twice weekly by replacing half the medium with fresh medium. After 12–14 days in vitro, cells underwent real-time oxygen consumption analysis with the Seahorse XFe24 machine and the MitoStress test kit. Primary astrocyte cultures were prepared from cerebral cortices of postnatal day 1–2 C57BL/6 pups. In brief, dissociated cortical cells were suspended in DMEM/F12 50/50 (Life Technologies, 11320-033) containing 25 mM glucose, 4 mM glutamine, 1 mM sodium pyruvate and 10% FBS, and plated on uncoated 75-cm^2^ flasks at a density of 1.5 × 10^5^ cells per cm^2^. Monolayers of astrocytes were obtained 12–14 days after plating. Cultures were gently shaken, and floating cells (microglia) were collected and removed, resulting in a more than 95% pure culture of astrocytes. Astrocytes were dissociated by trypsinization and then reseeded at 4 × 10^4^ cells per well in a XF24-well cell culture microplate, and cells underwent real-time oxygen consumption analysis with the Seahorse XFe24 machine and the MitoStress test kit.

### oAβ and oTau preparation

Oligomeric amyloid-beta was prepared as previously described (*5*). Amyloid-β_42_ was obtained from rPeptide (Cat. No. A-1163-2). To prepare amyloid-β 42 in oligomeric form (oAß), HFIP-prepared amyloid-β42 was resuspended in dimethyl sulfoxide (DMSO; 0.1 mg in 10 μl) followed by 1:10 dilution in Ham’s F12 culture medium (Mediatech) at 4°C for 24 h before use. This stock solution of 222 μM (molarity based on original amyloid-β_42_ monomer concentration) was then diluted in 1X PBS pH 7.4 for cell treatment experiments. Oligomeric Tau was prepared as previously described (*6*). Tau oligomers were prepared by dissolving 5 µM human recombinant tau from rPeptide (2N4R isoform; Cat. No. T-1001-2) in MES buffer (4-l pm morpholineethanesulfonic acid hydrate, pH 6.5), then mixed with 10 µM DTT (Sigma-Aldrich Corp. St Louis, MO), and incubated for 10 min at 55°C. Subsequently, 5 µM heparin (H19, Thermo Fisher Scientific, Waltham, MA) was added to the solution to induce aggregation and the solution was placed on the shaking incubator with a speed of 1000 rpm for 4 h at 37°C. Stock concentration of tau oligomers (oTau) was 5 µM and was diluted for cell treatment experiments. Vehicle consisted of DMSO in 1:10 Ham’s: F12 media for oAß and MES buffer, DTT, and heparin for oTau. Final concentrations in culture assays were 100 nM for oAß and 100 nM for oTau.

### Human iPSC-derived astrocyte induction and maturation

Cortical-like astrocytes were generated from the human iPSC line WTC11 as previously described (*7*). Briefly, on day 0 of differentiation, iPSCs were dissociated into small aggregates and transferred to untreated tissue culture flasks with SMAD inhibitors SB431542 (Stemcell Tech, 72234) and DMH1 (Tocris, 73634). On day 7 when embryoid bodies began to show rosette clusters, spheroids were transferred to Matrigel (Corning, 354230) -coated tissue culture plates without SMAD inhibitors. On day 14, rosette clusters were mechanically removed and transferred to tissue culture flasks with FGFb (Peprotech, 100-18B). On day 20, spheroids were triturated into a single cell suspension and transferred to a new untreated cell culture flask. From Day 28 to 180, spheroid aggregates were maintained in suspension with EGF (Peprotech, 100-15) and FGFb. Media was changed every 4-5 days. Spheroid aggregates were triturated with Accutase (Gibco,A1110501) every 7-10 days and transferred to new untreated tissue culture flasks. For the maturation of astrocytes, spheroids were triturated and plated onto Matrigel-coated 24-well plates with BMP4 (Peprotech, 120-05ET) and CNTF (Peprotech, 450-13) for 1 week.

### Mass labeling and LC/MS of KP and NAD+ metabolites

Isotope labeling of cells was performed as previously described (*8*). [U-13C] Trpytophan (Cat no. CLM-4290-H-PK, Cambridge Isotope Laboratories, Cambridge, MA), or unlabeled L-Tryptophan (Cat no. T0254, Sigma-Aldrich, St. Louis, MO) were supplemented to customized RPMI media lacking L-Tryptophan (Cat no. 72400120, Customized RPMI 1640 Medium + GlutaMAX supplement + HEPES, ThermoFisher, Pittsburgh, PA). For steady state labeling of metabolites [U-13C] Tryptophan (0.025 mM), labeled medium was replaced every day, and 2h before extracting metabolites. Metabolism was quenched by rapidly cooling cells on dry ice and cells were washed with 1x PBS twice before metabolites were extracted by aspirating media and immediately adding 1 mL -80°C 80:20 methanol: water. After 20 min of incubation on dry ice, the resulting mixture was scraped, collected into a centrifuge tube, and centrifuged at 10,000g for 5 min at 4°C.

LC/MS was performed as previously described (*9, 10*). In brief, the LC–MS method involved hydrophilic interaction chromatography (HILIC) coupled to the Q Exactive PLUS mass spectrometer (Thermo Scientific). The LC separation was performed on a XBridge BEH Amide column (150 mm × 2.1 mm, 2.5 μm particle size, Waters, Milford, MA). Solvent A is 95%: 5% H_2_O: acetonitrile with 20 mM ammonium bicarbonate, and solvent B is acetonitrile. The gradient was 0 min, 85% B; 2 min, 85% B; 3 min, 80% B; 5 min, 80% B; 6 min, 75% B; 7 min, 75% B; 8 min, 70% B; 9 min, 70% B; 10 min, 50% B; 12 min, 50% B; 13 min, 25% B; 16 min, 25% B; 18 min, 0% B; 23 min, 0% B; 24 min, 85% B; 30 min, 85% B. Other LC parameters are: flow rate 150 µl/min, column temperature 25 °C, injection volume 5 μL. The mass spectrometer was operated in positive ion mode for the detection of NAD, and negative ion mode for the KP metabolites (Trp, Kyn, 3-HANA, KYNA, QA). Other MS parameters are: resolution of 140,000 at m/z 200, automatic gain control (AGC) target at 3e6, maximum injection time of 30 ms and scan range of m/z 75-1000. All isotope-labeling patterns were corrected for natural abundance.

### Quantitative immunoblotting

Quantitative immunoblotting was carried out as previously described (*11*). Mouse anti-β-actin (1:10,000; Sigma-Aldrich) was used as an internal loading control. Imaging of blots was carried out using a Licor CLX-1306 with fluorescent secondary antibodies. Analysis was carried out in Image Studio Lite and signal intensities were normalized to loading controls when applicable. Antibodies used were as follows: anti-mouse aryl-hydrocarbon receptor (ThermoFisher Scientific, MA1-513, 1:500) and anti-β-actin antibody (Millipore Sigma, A5441, 1:10,000). Co-immunoprecipitation was performed using the Dynabeads Protein A Immunoprecipitation Kit (TheromFisher Scientific, 10006D) according to manufacturer’s instructions. In brief, 5 μg of ARNT antibody (Novus Biologicals, NB100-124) were incubated with 50 μL Dynabeads (ThermoFisher Scientific, 10001D). Antibody-bound beads were then incubated with 50 µg astrocyte cell lysates and allowed to bind antigen for 10 minutes at room temperature. Antibody-antigen complex was then eluted with manufacturer provided elution buffer and denatured with pre-mixed NuPAGE™ LDS Sample Buffer and NuPAGE Sample Reducing Agent. Samples were heated at 95° C for 5 minutes before loading on the gel.

### Immunocytochemistry

Mouse astrocytes were plated at a confluency of 1×10^5^ cells per well in a 12-well plate on top of coverslips coated with poly-L-lysine for 24h. Cells were then fixed in 4% PFA, washed 3 x with PBS with 0.1% Tween 20, and then blocked for 1 hour with 1% BSA and 10% NGS with 0.1% Tween 20. Coverslips were incubated with antibodies to GFAP (ThermoFisher Scientific, 13-0300, 1:1000) and to aryl-hydrocarbon receptor (ThermoFisher Scientific, MA1-513, 1:500). Z-stack images of GFAP+ cells spanning 15 μm were captured using a Keyence Fluorescent BZX-700 microscope. Coverslips were washed, stained with DAPI, and coverslipped as before.

### qRT-PCR for AhR and Hif-1α targets

RNA was isolated using Invitrogen TRIzol and purified with Invitrogen Purelink RNA or with a chloroform– phenol protocol as previously described (*5*). cDNA was generated from 300 ng of RNA by using the Applied Biosystems High Capacity RT kit. Quantitative real-time PCR (qRT–PCR) was performed on a QuantStudio6 using Hif-1α and AhR RT2 Profiler (Qiagen LLC, 330231) 384 plates with SYBR green qPCR master mix (Qiagen LLC, 330523). Relative expression was measured using the ΔΔCt method with beta-actin.

### LC/MS metabolomics of cultured cells

Metabolites were extracted from isolated astrocytes and neurons in a -80°C 80:20 methanol:water solution in a volume of 1.5 ml per 1×10^6^ cells, vortexed, incubated on dry ice for 10 min, and centrifuged at 16,000 g for 20 min, and the supernatant was assayed by LC-MS analysis. Extracts were analyzed within 24 hr by liquid chromatography coupled to a mass spectrometer (LC-MS). The LC-MS method involved hydrophilic interaction chromatography (HILIC) coupled to the Q Exactive PLUS mass spectrometer (Thermo Scientific) (*12*). The LC separation was performed on an XBridge BEH Amide column (150 mm 3 2.1 mm, 2.5 mm particle size, Waters, Milford, MA). Solvent A was 95%: 5% H2O: acetonitrile with 20 mM ammonium bicarbonate, and solvent B was acetonitrile. The gradient was 0 min, 85% B; 2 min, 85% B; 3 min, 80% B; 5 min, 80% B; 6 min, 75% B; 7 min, 75% B; 8 min, 70% B; 9 min, 70% B; 10 min, 50% B; 12 min, 50% B; 13 min, 25% B; 16 min, 25% B; 18 min, 0% B; 23 min, 0% B; 24 min, 85% B; and 30 min, 85% B. Other LC parameters are: flow rate 150 ml/min, column temperature 25°ΔC, and injection volume 10 mL and autosampler temperature was 5°ΔC. The mass spectrometer was operated in both negative and positive ion mode for the detection of metabolites. Other MS parameters were: resolution of 140,000 at m/z 200, automatic gain control (AGC) target at 3e6, maximum injection time of 30 ms and scan range of m/z 75-1000. Raw LC/MS data were converted to mzXML format using the command line “msconvert” utility (*13*). Data were obtained with MAVEN software (*14, 15*).

For identification of hexose phosphates and glycolytic intermediates, capillary electrophoresis mass spectroscopy was used as previously described (*16*). In brief, adherent cells on dishes were washed with 5% mannitol aqueous solution at room temperature. The cells were immersed in 400 μl methanol for 30 s, and 275 μl of the Internal Standard Solution (10 μM, Solution ID: H3304-1002, Human Metabolome Technologies) for 30 s. The extraction liquid was centrifuged at 2,300*g* for 5 min at 4 °C. The supernatant (400 μl) was centrifugally filtered at 9,100*g* for 4 h at 4 °C through a 5-kDa cut-off filter (Millipore) to remove proteins, and then the filtrate was lyophilized and suspended in 25 μl Milli-Q water. The metabolite suspension was analysed by capillary electrophoresis time of flight mass spectrometry (CE-TOF/MS) using an Agilent capillary electrophoresis (CE) system equipped with an Agilent 6210 TOFMS, an 1100 isocratic high-performance liquid chromatography pump, a G1603A CE-MS adaptor kit and a G1607A CE-electrospray ionization-mass spectrometry (ESI-MS) sprayer kit (Agilent Technologies). The system was controlled using G2201AA ChemStation software v.B.03.01 for CE (Agilent).

Metabolomics data analysis was carried out using MetaboAnalyst version 5.0 (*13*). Metabolomic data were log-transformed and scaled according to the auto-scaling feature (mean-centered and divided by the standard deviation of each variable). Metabolites that were significantly different by ANOVA (with FDR adjusted *P* value, or *q* < 0.05) were subjected to hierarchical clustering analysis using the Euclidian distance measure and Ward clustering algorithm. Differentially expressed metabolites by volcano plot underwent pathway7-based enrichment analysis using 84 metabolite sets based on KEGG human metabolic pathways in MetaboAnalyst 5.0.

### LC/MS metabolomics of brain tissues

Frozen brain tissues were weighed, ground with a liquid nitrogen in a cryomill (Retsch) at 25 Hz for 45 seconds, before extracting tissues 40:40:20 acetonitrile: methanol: water +0.5% FA +15% NH4HCO3 (*17*) with a volume of 40 L solvent per 1 mg of tissue, vortexed for 15 seconds, and incubated on dry ice for 10 minutes. Brain samples were then centrifuged at 16,000 g for 30 minutes. The supernatants were transferred to new Eppendorf tubes and then centrifuged again at 16,000 g for 25 minutes to remove any residual debris before analysis.

Extracts were analyzed within 24 hours by liquid chromatography coupled to a mass spectrometer (LC-MS). The LC–MS method was based on hydrophilic interaction chromatography (HILIC) coupled to the Orbitrap Exploris 240 mass spectrometer (Thermo Scientific) (*12*). The LC separation was performed on a XBridge BEH Amide column (2.1 x 150 mm, 3.5 m particle size, Waters, Milford, MA). Solvent A is 95%: 5% H_2_O: acetonitrile with 20 mM ammonium acetate and 20 mM ammonium hydroxide, and solvent B is 90%: 10% acetonitrile: H2O with 20 mM ammonium acetate and 20 mM ammonium hydroxide. The gradient was 0 min, 90% B; 2 min, 90% B; 3 min, 75% B; 5 min, 75% B; 6 min, 75% B; 7 min, 75% B; 8 min, 70% B; 9 min, 70% B; 10 min, 50% B; 12 min, 50% B; 13 min, 25% B; 14min, 25% B; 16 min, 0% B; 18 min, 0% B; 20 min, 0% B; 21 min, 90% B; 25 min, 90% B. The following parameters were maintained during the LC analysis: flow rate 150 ml/min, column temperature 25 °C, injection volume 5 µl and autosampler temperature was 5°C. For the detection of metabolites, the mass spectrometer was operated in both negative and positive ion mode. The following parameters were maintained during the MS analysis: resolution of 180,000 at m/z 200, automatic gain control (AGC) target at 3e6, maximum injection time of 30 ms and scan range of m/z 70-1000. Raw LC/MS data were converted to mzXML format using the command line “msconvert” utility (*18*). Data were analyzed via the EL-MAVEN software version 12.

### OCR and ECAR

Cells were counted and plated at 1 × 10^5^ (mouse astrocytes or neurons) cells per well in a Seahorse XF24 Cell Culture Microplate for all experiments (Agilent). Cells were then treated with indicated inhibitors or agonists in each experiment for 20 h. Cells were washed twice with Agilent Seahorse XF medium (Agilent) supplemented with 1 mM pyruvate, 2 mM L-glutamine and 10 mM D-glucose; a final volume of 525 μl was placed in each well. Cells were then incubated in a 0% CO2 chamber at 37 °C for 1 h before being placed into a Seahorse XFe24 Analyzer (Agilent). For OCR and ECAR mitostress test experiments, cells were treated with 1 μM oligomycin, 2 μM FCCP and 0.5 μM rotenone or antimycin (indicated by three black arrows in representative OCR traces). All Seahorse experiments were repeated at least three times. All OCR and ECAR data were normalized to cell number per well using CyQUANT (Thermo Fisher Scientific).

### Barnes maze

The Barnes maze protocol was adopted from a previous study (*19*) with minor modifications. The maze was made from a circular, 8-mm thick, white PVC slab with a diameter of 36 inches. Twenty holes with a diameter of 3 inches were made on the perimeter at a distance of 1 inch from the edge. This circular platform was then mounted on top of a rotating stool, 30 inches above the ground. The escape cage was made by using a mouse cage and assembling a platform and ramp 2 inches below the surface of the maze. The outside of the walls of the cage were covered with black tape so as to prevent light for entering the escape cage. The maze was placed in the centre of a dedicated room and two 120-W lights were placed on the edges of the room facing towards the maze to provide an aversive stimulus for the mice. Eight simple colored-paper shapes (squares, rectangles and circles) were mounted on the walls of the room as visual cues. After testing each mouse, the maze was cleaned with 70% ethanol and rotated clockwise after every mouse to avoid intra-maze odor or visual cues. All sessions were recorded using a JVC Everio HD camcorder GZ-E200 and analyzed with Kinovea video tracking software. The mice interacted with the Barnes maze in three phases: habituation (day 1), training (days 2–3) and probe (day 4). Before starting each experiment, mice were acclimated to the testing room for 1 h. Then all mice from one cage (n = 4 or 5) were placed in individual holding cages, where they remained until the end of their testing sessions each day. On habituation day, the mice were placed in the center of the maze within a vertically oriented black PVC pipe 4 inches in diameter and 7 inches in height for 15 s. The mice were then guided slowly to the hole that led to the escape cage over the course of 10–15 s. The mice were given 3 min to independently enter the target hole, and if they did not, they were nudged with the PVC pipe to enter. The 120-W lights were then shut off and mice were allowed to rest in the escape cage for 2 min. The training phase occurred 24 h after the habituation phase and was split across 2 days (days 2 and 3), with 3 trials on the first day and 2 trials on the second day. During each trial, the mice were placed in the center of the maze within the PVC pipe for 15 s and afterwards were allowed 3 min to explore the maze. If mice found and entered the target hole before 3 min passed, the lights were shut off and the training trial ended. Mice were allowed to rest in the escape cage for 2 min. If at the end of the three minutes the mice had not entered the target hole, they were nudged with the PVC pipe. A total of five trials were conducted. During each trial, latency (time) to enter the target hole as well as distance travelled were recorded. The probe phase occurred 24 h after the training phase and was conducted on the last day (day 4). Mice were placed in the center of the maze within the PVC pipe for 15 s and afterwards were allowed 3 min to explore the maze. The probe session ended whenever the mouse entered the target hole or if 3 min had passed. During the probe phase, measures of time spent per quadrant, latency to enter the target hole and distance travelled were recorded. Behavioral testing was performed and evaluated by researchers blinded to treatment.

### Novel object recognition (NOR) task

The NOR task is based on the ability of mice to show preference for novel versus familiar objects when allowed to explore freely and was adopted from a previous study (*20*). NOR was performed during the light cycle. Mice were individually habituated to an open arena (50 cm × 50 cm, dim light, 24°C) on day 1 for approximately 5 min. During the subsequent training session, 2 identical objects were placed into the arena, and exploratory behavior was monitored for 10 min. On day 2, mice were placed back into the same arena, in which one of the objects used during training was replaced by a novel object of similar dimensions but a different shape/color, and exploratory behavior was monitored for 5 min. Digital video tracking (using an infrared camera and vplsi Viewpoint software) of body movements and head position was used to quantify locomotor and exploratory activities around the objects (2-cm zone around the objects). Exploration behavior was assessed by calculating DI, the ratio of number of interactions exploring the new object divided by the old object in the testing phrase normalized to the number of interactions spent observing each respective old object the day prior. A DI of approximately 1 indicates association with correct training and no object preference; a significant increase in DI is characteristic of recognition of the novel object. Behavioral testing was performed and evaluated by experimenters blinded to treatment or genotype.

### Morris water maze

The Morris water maze (MWM) was conducted as previously described (*21*). MWM was originally designed to test spatial reference memory in rats by observing and recording escape latency, distance moved, and velocity during the search for a hidden escape platform in a large pool (Morris 1984). For our test, we used a large water tank (178 cm in diameter) filled with water at a temperature of 22.0 ± 1.5◦C with a circular platform (17 cm in diameter) placed about 1 cm below the water surface and approximately 50 cm away from the wall. Nontoxic tempera paints (Elmers, Westerville, OH) were used to make the water opaque. The water tank was completely surrounded by privacy blinds with at least 4 visual cues attached to the blinds. Four different shapes including a star shape, circle, rectangle, and diamond, each with approximately 6 square feet of surface area were used as visual cues. The visual cues were located approximately 150 cm from the center of the tank. The water tank arena was monitored by an overhead video system that allowed Ethovision to track the mice. During hidden platform training, a platform was positioned in one quadrant of the tank. Mice were released from pseudorandomized drop locations and given 90 sec to find the platform. The distance to the platform was generally the same within a day. The trial either ended when the mice rested on the platform for 10 sec or until the trial duration expired. If mice failed to find the submerged hidden platform during that time, they were guided to it. Mice underwent 4 trials of training each day (30-min ITIs) for four consecutive days. Upon completion of the hidden platform training, the platform was removed and a 30-sec probe trial was conducted. Successful learning of MWM was determined by the gradual decrease in escape latency and discriminative quadrant exploration during the probe trial. For analysis, data were averaged per day. After the probe trial, mice were given visible platform training to ensure that no gross sensorimotor or visual deficits were present. During the visible platform training, the platform was marked with a black-and-white ping-pong ball attached to a 10-cm wooden stick. No mice were excluded based on our standard exclusion criteria in this task: excessive thigmotaxis, obvious visual impairment, excessive corkscrew swimming pattern, and obvious sensorimotor dysfunction. The water was frequently changed and the tank disinfected. Behavioral testing was performed and evaluated by researchers blinded to genotype.

### Measurement of Aß42 peptides

Measurement of total Aß42 was performed in guanidine brain lysates as previously described (*22*).

### Electrophysiology

To measure the cellular mechanism of learning and memory, we used a modified protocol previously described (*23*). In brief, mice were euthanized by cervical dislocation, and the hippocampus was rapidly dissected into ice-cold (4 °C) artificial cerebrospinal fluid (ACSF), saturated with carbogen (95% O_2_/5% CO_2_). ACSF consisted of (in mM): 124 NaCl, 4.9 KCl, 24.6 NaHCO_3_, 1.20 KH_2_PO_4_, 2.0 CaCl_2_, 2.0 MgSO_4_ and 10.0 glucose, pH 7.4. Transverse hippocampal slices (350 μm thick) were prepared from the dorsal area of the hippocampus with the McIlwain tissue chopper and transferred to a recovery chamber for at least 1.5 h with oxygenated ACSF at room temperature before being placed into a submerged-type chamber in which they were kept at 32 °C and continuously perfused with ACSF at a flow-rate of 1.5 ml/min. Slices were carefully positioned on a R6501A multi-electrode array (Alpha MED Scientific), with electrodes arrayed in an 8 × 2 matrix with interpolar distance of 150 μm; each matrix measured 50 μm × 50 μm. After a 30-min incubation, the fEPSPs in CA1 were recorded by stimulating downstream electrodes in the CA1 and CA3 regions along the Schaffer collateral pathway. Signals were acquired using the MED64 System (AlphaMED Sciences, Panasonic). The time course of the fEPSP was calculated as the descending slope function for all experiments. Input/output curves were established by applying increasing stimulus currents to the pathway from 10 μA to 90 μA (in 5-μA increments) and recording evoked responses. After input/output curves had been established, the stimulation strength was adjusted to elicit a fEPSP slope at 35% maximal value, which was maintained throughout the experiment. During baseline recording, a single response was evoked at a 30-sec interval for at least 20 min. To induce a strong form of long-term potentiation, three episodes of theta-burst stimulation (TBS) were used, each TBS consisting of 10 bursts of 4 stimuli at 100 Hz separated by 200 ms (double pulse-width), followed by recording evoked responses beginning 1 min after the induction of long-term potentiation and continuing every 30 sec until the end of the experiments. Experiments using control and transgenic mice were interleaved with each other. The mean baseline fEPSP value was calculated and percentage change from baseline after the TBS was analyzed for long-term potentiation.

APP/PS, 5XFAD, and PS19 mice were treated with vehicle or PF068 at 15 mg/kg/day for one month, euthanized and hippocampal slices were assessed by electrophysiology. The MCT1/2 inhibitor AR-C155858 (*24*) (Tocris) was prepared fresh for each experiment and was diluted to a final concentration of 50 µM in ACSF and applied via the perfusion line during baseline recordings. For LTP experiments, all drugs were applied 15 min after the start of recording and 45 min prior to TBS. MCTi was left on the slice for the entire recording interval. All drug experiments were interleaved with vehicle controls.

### Carotid artery catheterization

Carotid artery catheters using polyethylene tubing (PE10, Braintree Scientific Inc.) were prepared as previously described (*25*). Anesthesia of mice was induced by inhalation of 5% isoflurane and maintained by inhalation of 2% isoflurane. The surgical site was prepared with skin disinfectant (betadine followed by alcohol wipes x3 followed by a final betadine wash). An incision was made above the jugular vein on the neck (approximately 7mm in length), at about 5 mm to the right of midline. The carotid artery was located and blunt dissected from surrounding tissue with bent tweezers. A pair of 6-0 silk sutures was placed below the artery and the catheter, which was filled with sterile filtered 10U Heparin/saline, was then inserted into the artery through the incision. A peristaltic pump (Harvard Apparatus, HA1100WD) connected to the catheter then began administering 0.8M U-^13^C-glucose at a steady rate of 4μl/min over the course of 2 hours. After 2 hours, mice were euthanized by CO_2_ asphyxiation followed by cervical dislocation. Brains were harvested and immediately frozen on dry ice for sectioning and subsequent MALDI analysis.

### Matrix-Assisted Laser Desorption/Ionization (MALDI)

Matrix-assisted laser desorption/ionization (MALDI) imaging analyses were performed as described previously (*26*). Briefly, thin sections (8 µm) of the brain were prepared with a cryomicrotome (CM3050, Leica Microsystems, Tokyo, Japan). Sections were attached onto indium tin oxide–coated glass slides (Bruker Daltonics GmbH, Leipzig, Germany) and were coated with 9-aminoacridine as the matrix (10 mg/mL, dissolved in 80% ethanol) by manually spraying with an artistic-brush (Procon Boy FWA Platinum, Mr. Hobby, Tokyo, Japan). The matrix was simultaneously applied to multiple sections to maintain consistent analyte extraction and co-crystallization conditions. MALDI imaging was performed using an Ultraflextreme MALDI-TOF/TOF mass spectrometer equipped with an Nd:YAG laser. Data were acquired in the negative reflectron mode with raster scanning at a pitch distance of 50 µm. Each spectrum was the result of 300 laser shots at each data point. For TOF/TOF measurement, signals between *m/z* 50 and 1000 were collected. Image reconstruction for both procedures was performed using FlexImaging 4.1 software (Bruker Daltonics). Molecular identification was based on accurate *m/z* value provided by FT-ICR-MS data and previous studies (*26, 27*).

### Human brain tissues

Postmortem brain material was obtained from the Arizona Study of Aging and Neurodegenerative Disorders and Brain and Body Donation Program (*28*). AD was defined as intermediate or high probability that dementia was due to AD, according to NIA-Reagan criteria (*29*). For LC-MS experiments, patients consisted of 4 groups stratified by Braak stage (*30*), including non-demented Braak I–II - zero to sparse plaques (n=12), AD Braak III–IV-demented with moderate plaques (n=12), and AD Braak V–VI - demented with frequent plaques (n=12). We excluded cases with clinicopathologic evidence of Parkinson’s disease (defined as having 2 of the 3 cardinal clinical signs of resting tremor, muscular rigidity and bradykinesia, along with pigmented neuron loss and Lewy bodies in the substantia nigra), dementia with Lewy bodies, progressive supranuclear palsy, motor neuron disease (including amyotrophic lateral sclerosis, primary lateral sclerosis and motor neuron disease associated with frontotemporal lobar degeneration), corticobasal degeneration (defined by the classic H&E histopathology of achromatic, swollen neurons in the cerebral cortex and indistinct inclusions within pigmented neurons of the substantia nigra as well as abnormal, phosphorylated tau or Gallyas-positive astrocytic plaques in the cerebral cortex), Pick’s disease (defined as clinical dementia with tau or silver-stain positive Pick bodies within neurons of the cerebral cortex, hippocampus and/or basal ganglia), multiple system atrophy (defined by atrophy and gliosis of the cerebellar folia, basal pons, substantia nigra and/or striatum, as well as alpha-synuclein or silver-positive glial and neuronal cytoplasmic inclusions), and Huntington’s disease (defined by the characteristic trinucleotide repeat expansion in the gene for huntingtin). Brains from subjects with cancer, sepsis, or stroke were also excluded. The post-mortem interval was less than 8 hours. Cases were balanced for sex in each group and all groups were matched for age at death. Demographic information is listed in **Supplementary Table 1**.

### Human control and LOAD patient iPSCs and differentiation to astrocytes and neurons

Patient and control individuals were diagnosed at the Shiley-Marcos UCSD Alzheimer’s Disease center at the University of California San Diego. Subjects were anonymized from researchers and dermal human fibroblasts were donated with informed consent and strict adherence to legal and ethical guidelines. Fibroblasts were reprogrammed to iPSCs by the Salk Stem Cell Core facility using commercial, non-integrating Sendai virus Yamanaka factors (CytoTune™-iPS 2.0 Sendai Reprogramming Kit; ThermoFisher Cat # A165167) per manufacturer’s recommendation. All fibroblasts and iPSCs were screened for mycoplasma monthly via MycoAlert™ PLUS Mycoplasma Detection Kit (Lonza Cat # 75860-362). iPSCs were grown in medium-sized colonies on Matrigel (BD Biosciences cat#: 354230) at a final concentration of 1 mg per 6-well plate and propagated in StemMACS™ iPS-Brew XF (Miltenyi cat # 130-104-368) with daily medium replacement. hAstrocytes were generated as previously described (*31*). hNeurons were generated via direct conversion through expression of doxycycline-inducible NGN2 and ASCL1 as previously described (*32*).

### Co-cultures of isogenic hNeurons and hAstrocytes

Co-culture experiments were performed in Corning Transwell (12-well) plates purchased from Thermo Fisher Scientific (Cat # 07-200-156). hNeurons were passaged onto 12-well plates at a density of 3×10^5^ cells per well. Mature hAstrocytes were plated on transwell inserts at a density of 3×10^5^ cells per insert. All experiments were performed in triplicate. Cells were allowed to recover in separate plates overnight before being combined for co-culture. hAstrocyte inserts were combined with hNeuron plates and allowed to equilibrate for 48 hours in DMEM/F12 containing N2, B27, with 5% KnockOut Serum replacement (Gibco, Cat # 10828028). hAstrocyte transwells were then moved to a new plate and treated with either vehicle (DMSO) or 200 nM PF068 overnight in glucose free Advanced DMEM/F12 (Gibco, Cat # A2494301) supplemented with ^13^C-glucose at 17.5 mM (Cambridge isotopes). Media on hNeuron wells were also replaced with similar media without treatment and in low glucose (5.5mM) to equilibrate overnight. hAstrocyte transwells were then washed with PBS and reintroduced to hNeuron plates containing freshly applied Advanced DMEM/F12 for 4 hours, at which point media were harvested and filtered, and cells were lifted, washed, pelleted and snap frozen for metabolomic analysis or transcriptomics.

### RNA sequencing

RNA was extracted from pelleted cells via TRIzol reagent (Invitrogen, Cat # 15596026) per manufacturer recommendations followed by Zymo Direct-zol column purification kit (Zymo Research, Cat # R2062). Library preparation was performed in 2 equal batches by Salk’s Next Generation Sequencing Core. Pair-end sequencing was performed on an Illumina NovaSeq6000 sequencer at a targeted read depth of 20 M reads per sample. Illumina adapters were trimmed with TrimGalore (*33*). Remaining reads were aligned to GRCh38 with STAR (*34*). Gene expression values and annotations were obtained through RSEM (*35*). Differential expression was performed with DESeq2 (*36*). Motif analysis was performed using findMotifs.pl in Homer (*37*). To avoid statistical artifacts, findMotifs.pl was run in quadruplicate and with randomized gene lists.

### Statistics

Data are expressed as mean ± s.e.m. unless otherwise indicated. Statistical comparisons were made in the Prism software using a Student’s t-test (for 2 groups meeting the normal distribution criteria, according to the Shapiro–Wilk normality test), Mann–Whitney U-test (for 2 groups not meeting the normal distribution criteria) or ANOVA with Tukey’s multiple comparison test (for groups across variables, with multiple comparisons between groups). Data were subjected to Grubbs’ test to identify the presence or absence of outlier data points; no outliers were excluded in our analyses. For all tests, P < 0.05 was considered significant, except for targeted metabolomics in which FDR-adjusted *P* value, or *q* < 0.05.

---

**Supplementary Data Table 1:**
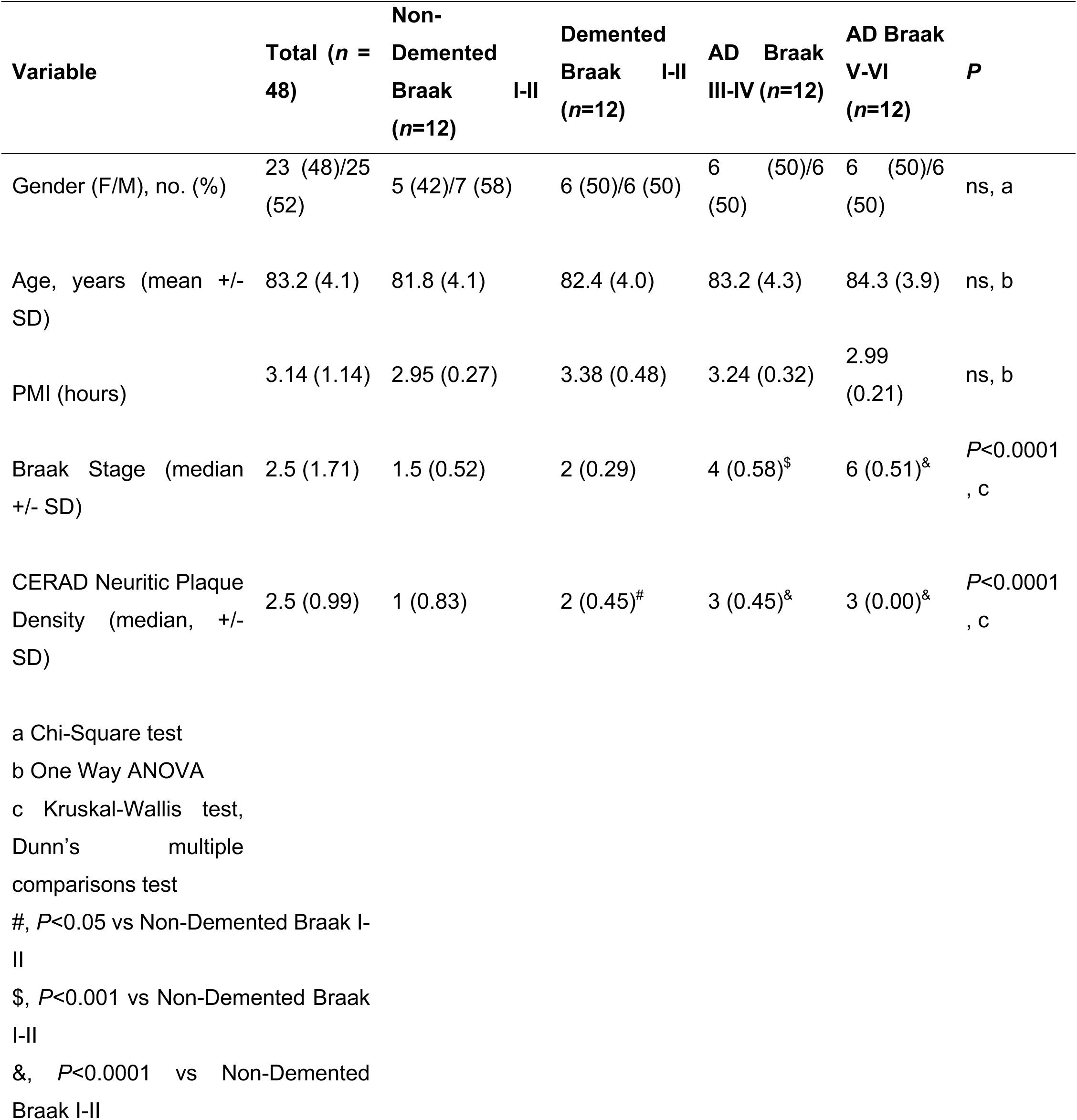
Demographic Characteristics of Study Participants.

**Supplementary Fig. 1.**
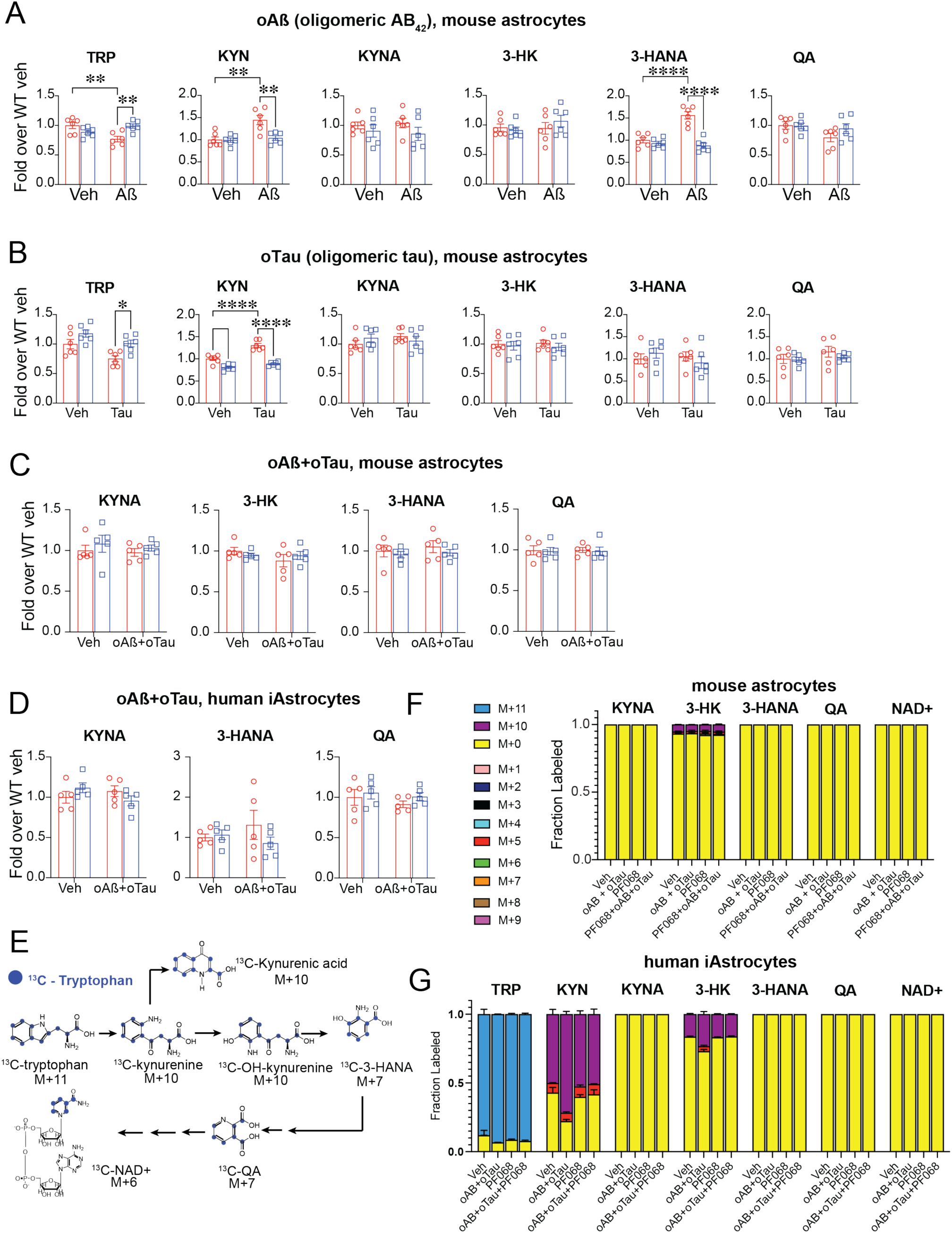
Mouse and human astrocytes upregulate IDO1 activity and KYN production in response to amyloid-beta and tau oligomers. Data are mean ± s.e.m. and analyzed using two-way ANOVA with Tukey post hoc tests: \**P*<0.05, \*\**P*<0.01, \*\*\**P*<0.001 and \*\*\*\**P*<0.0001. TRP = tryptophan, KYN = kynurenine, KYNA = kynurenic acid, 3-HK = 3-hydroxkynurenine, 3-HANA = 3-hydroxyanthralinic acid, QA = quinolinic acid **(A)** LC-MS quantification of kynurenine pathway (KP) metabolites in mouse astrocytes stimulated with oAβ (100 nM, 20h) +/- PF068 (100nM, 20h; n=6/group). **(B)** LC-MS quantification of KP metabolites in mouse astrocytes stimulated with oTau (100 nM, 20h) +/- PF068 (100nM, 20h; n=6/group). **(C)** LC-MS quantification of downstream KP metabolites in mouse astrocytes stimulated with oAβ+oTau (both at 100 nM, 20h) +/- PF068 (100nM, 20h; n=5/group). **(D)** LC-MS quantification of KP metabolites iAstrocytes +/- oAß+oTau (100 nM each, 20 h; n=5/group). **(E)** Diagram of isotope labeling of 13C Tryptophan and the KP. Mass labelled carbons are in purple. **(F)** Isotope tracing of ^13^C-tryptophan [M+11] for metabolites downstream of KYN in mouse astrocytes in response to oAβ+oTau (both at 100 nM, 20h) +/- PF068 (100nM, 20h; n=5/group) **(G)** Isotope tracing of ^13^C-tryptophan [M+11] to KYN [M+10] and KP metabolites in human iAstrocytes in response to oAβ+oTau (both at 100 nM, 20h) +/- PF068 (100nM, 20h; n=5/group).

**Supplementary Fig. 2.**
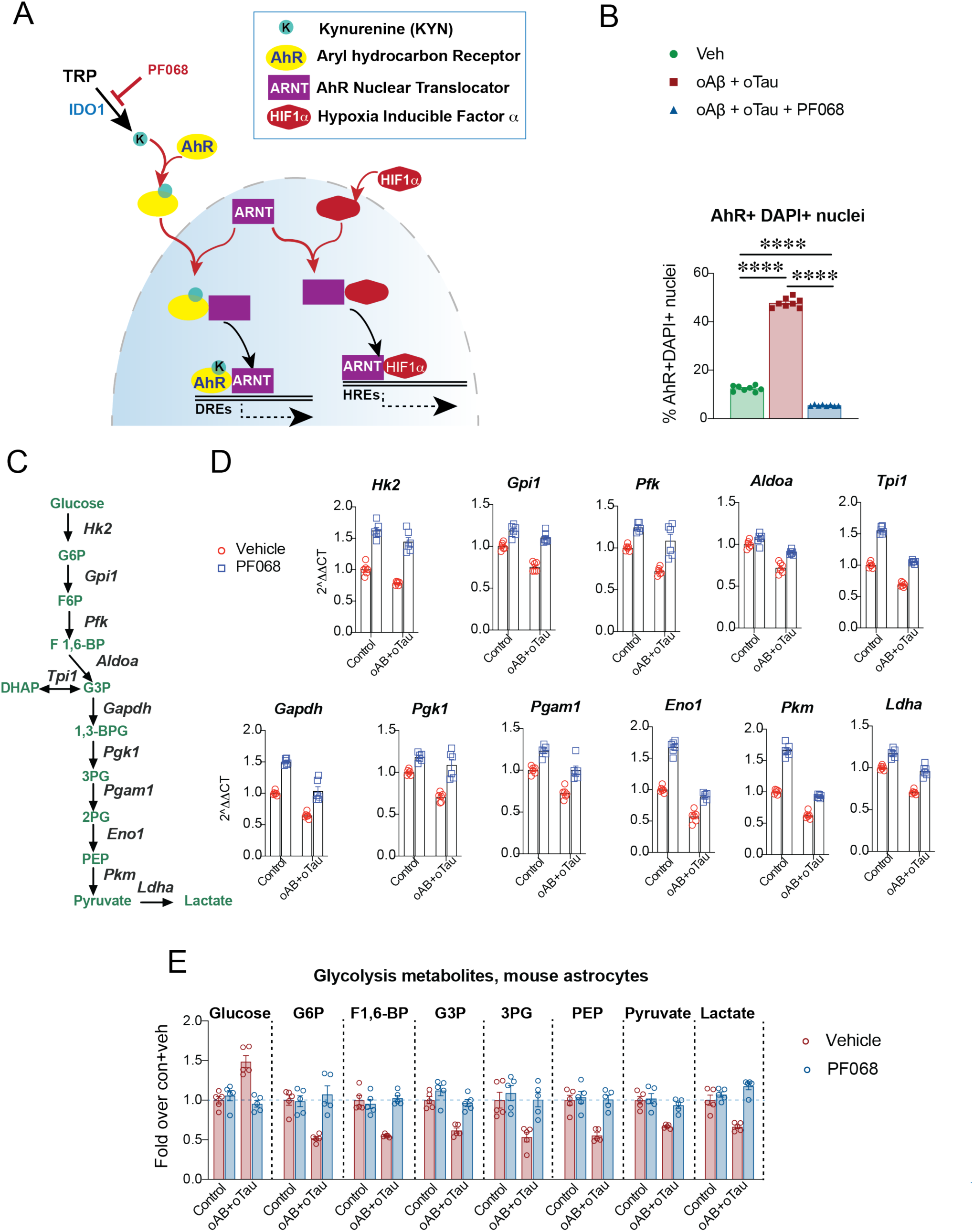
IDO1 inhibition and reduction of KYN-AhR nuclear translocation increase glycolytic gene expression and glycolytic and lactate intermediates. **(A)** Diagram of interaction between AhR:ARNT and HIF1α:ARNT. KYN is produced upon IDO1 activation and binds in the cytoplasm to AhR, whereupon the AhR-KYN complex translocates to the nucleus to bind ARNT and transcribe DRE genes. ARNT in the nucleus is limiting, and in the absence of AhR-KYN will bind instead to HIF1α. Inhibition of IDO1 activity will reduce transcription of AhR-regulated genes and conversely enhance transcription of HIF1α HRE genes, which include glycolytic genes. **(B)** Quantification of percent AhR+/DAPI+ nuclei in mouse astrocytes stimulated with veh or oAβ+oTau (100nM, 20h) +/- PF068 (100nM, 20h; n=8/group); \*\*\*\**P*<0.0001 by one-way ANOVA with Tukey’s post-hoc test. **(C)** Diagram of glycolysis pathway with genes encoding glycolytic enzymes in italics. **(D)** Quantification of transcript levels of glycolytic genes from **(C)** in mouse astrocytes stimulated with oAβ+oTau (100nM, 20h) +/- PF068 (100nM, 20h; n=5/group). **(E)** LC/MS quantification of glycolytic intermediates from mouse astrocytes stimulated with oAβ+oTau (100nM, 20h) +/- PF068 (100nM, 20h; n=5/group, 20h).

**Supplementary Fig. 3.**
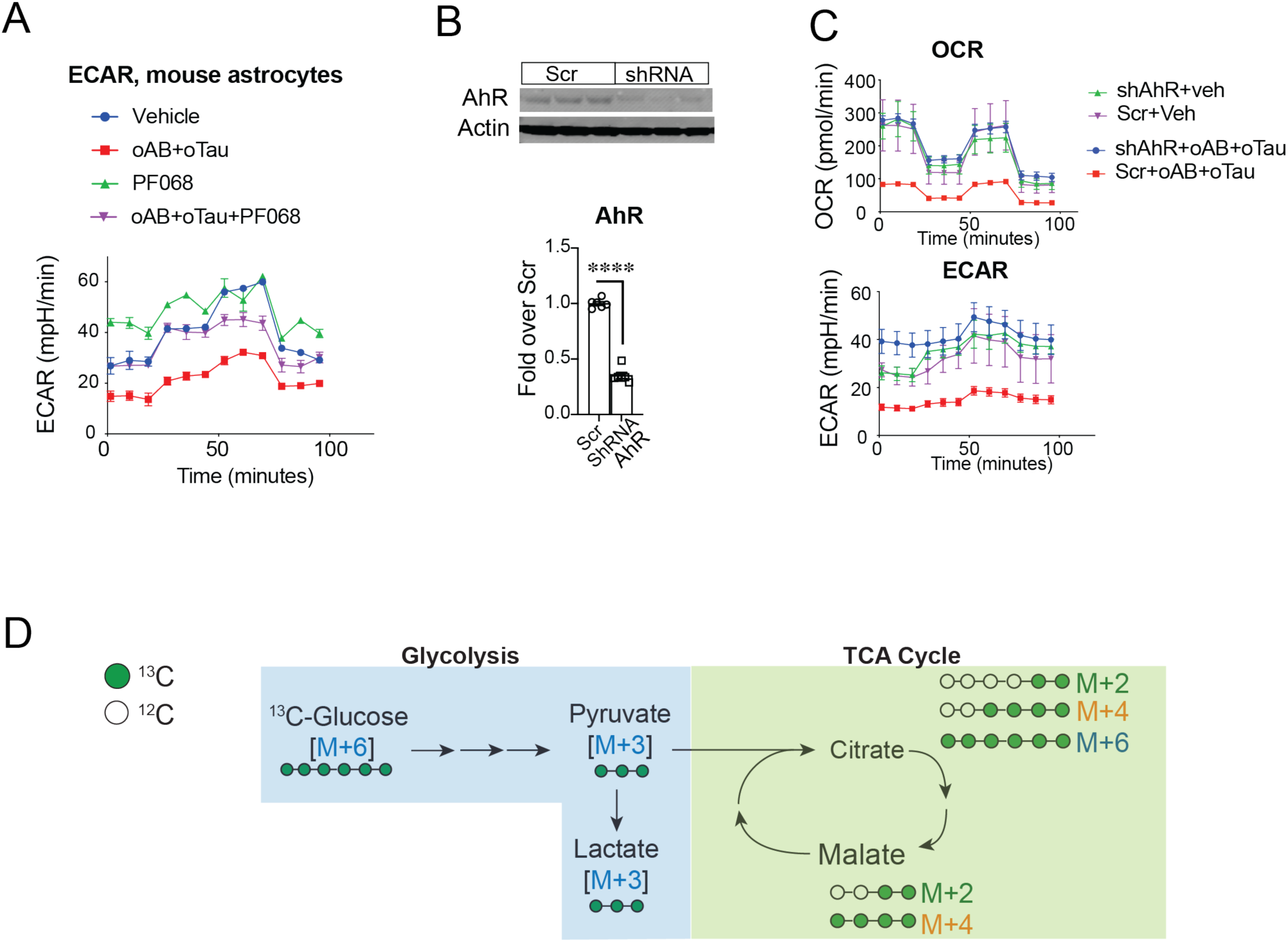
Inhibition of IDO1 in astrocytes. **(A)** Real time ECAR trace in mouse astrocytes stimulated with veh or oAβ+oTau (100nM, 20h) +/- PF068 (100nM, 20h; n = 5/group) **(B)** Representative immunoblot of AhR in mouse astrocytes transfected with shRNA to AhR, and quantification of AhR relative to actin, n=5/ group, \*\*\*\**P*<0.0001 by two-tailed unpaired *t*-test. **(C)** Real-time OCR and ECAR traces in mouse astrocytes transfected with shRNA to AhR and stimulated with veh or oAβ+oTau (100nM each, 20h; n = 5/group). **(D)** Diagram of isotope labeling of ^13^C-glucose and metabolism down the glycolytic pathway and TCA cycle.

**Supplementary Fig. 4.**
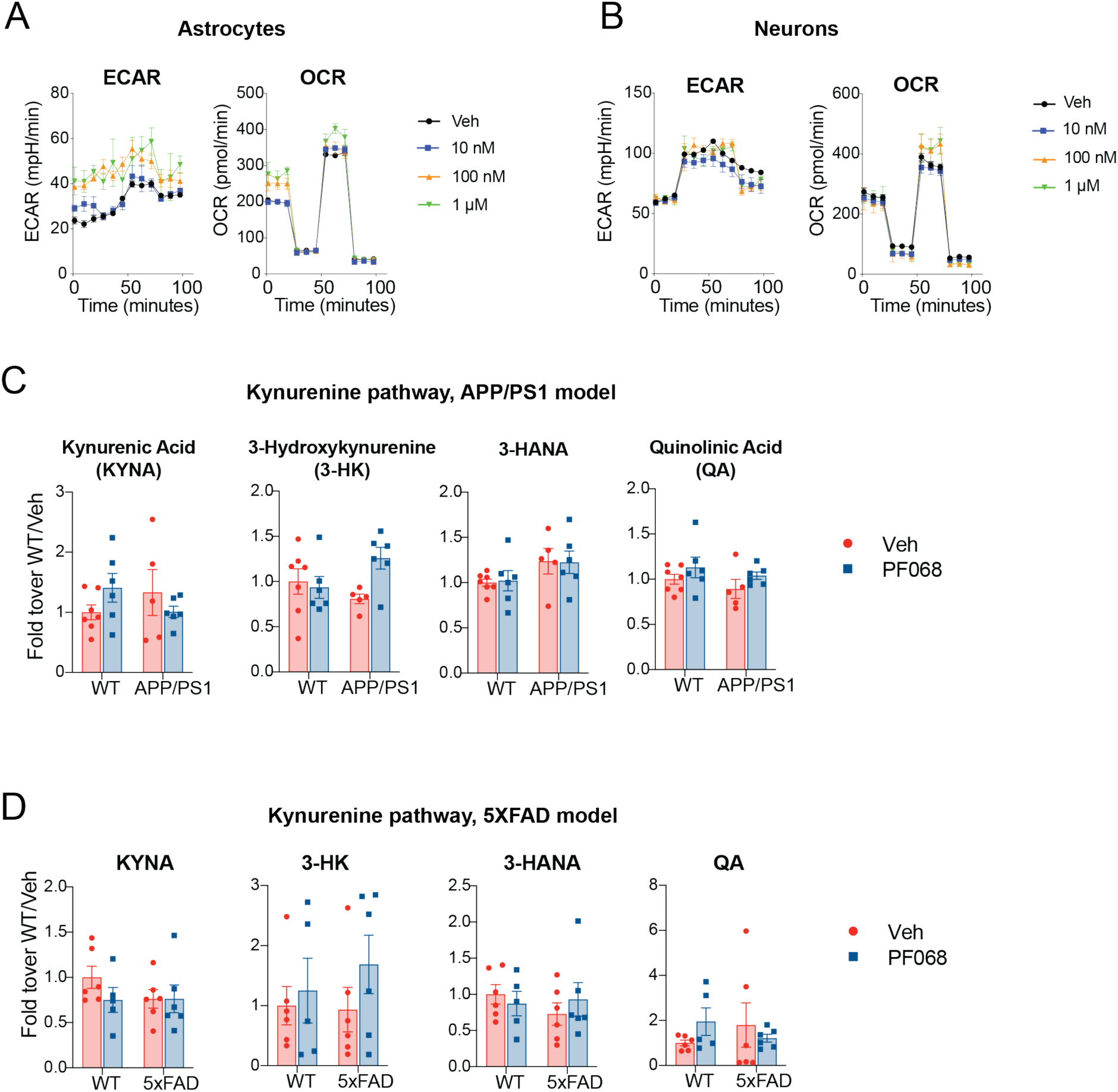
Effects of IDO1 inhibition on KP metabolites in mutant APP mice. Data are mean ± s.e.m. and are analyzed using two-way ANOVA with Tukey post hoc tests: KYNA = kynurenic acid, 3-HK = 3-hydroxkynurenine, 3-HANA = 3-hydroxyanthralinic acid, QA = quinolinic acid **(A)** Tracing of ECAR and OCR in mouse astrocytes treated with dose response of PF068 (n = 5 wells/group, 20h). **(B)** Tracings of ECAR and OCR in mouse neurons treated with dose response of PF068 (n = 9-10 wells/group, 20h). **(C)** LC/MS measurements of KP metabolites downstream of KYN in WT and APP/PS1 mice +/- PF068 at 15 mg/kg/day for one month (11-13 months old, n=5-7/group). **(D)** LC/MS measurements of KP metabolites downstream of KYN in WT 5XFAD mice +/- PF068 at 15 mg/kg/day for one month (6-7 months old, n=5-6/group).

**Supplementary Fig. 5.**
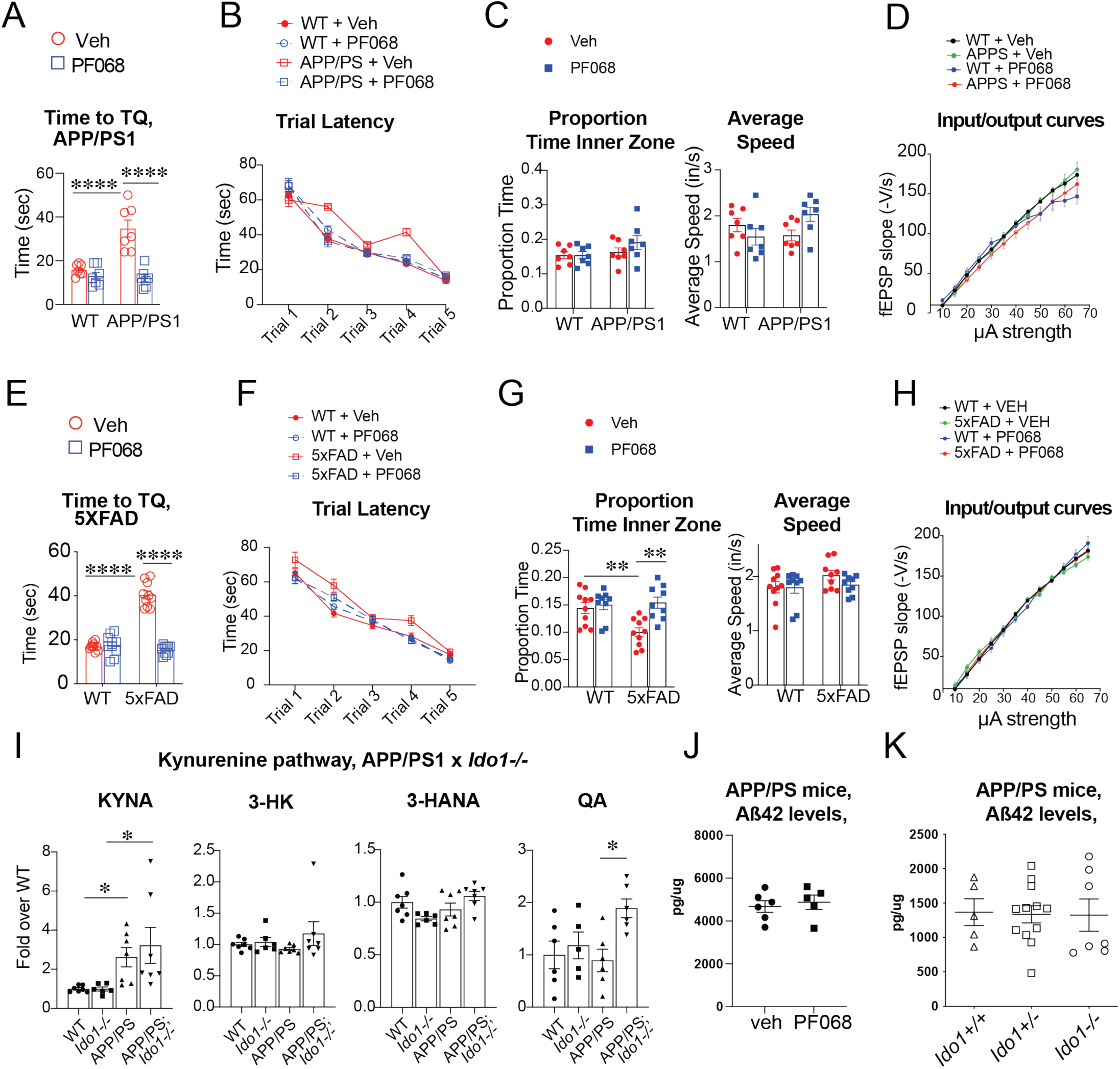
Effects of IDO1 inhibition on spatial memory in mutant APP mice. Data are mean ± s.e.m. and are analyzed using two-way ANOVA with Tukey post hoc tests: \**P*<0.05, \*\**P*<0.01, \*\*\*\**P*<0.0001. APP/PS1 mice and WT littermates (10-12 months old, n=7/group) and 5xFAD mice (5-6 months old, n=9-10/group) were treated with veh or PF068 at 15 mg/kg/day for one month and then behaviorally tested. **(A)** Time to TQ or primary latency (sec) to the target hole in the Barnes Maze task for APP/PS1 mice. **(B)** Primary latency in the Barnes maze for the five learning trials. **(C)** Percentage of time spent in the inner zone in the NOR task and average speed (inches per second). **(D)** Input/output curves as a measure of basal synaptic transmission in the CA1 region of the hippocampus (n = 8-9 slices, 4-5 mice/group). **(E)** Time to TQ or primary latency (sec) to the target hole in the Barnes Maze task for 5XFAD mice. **(F)** Primary latency in the Barnes maze for the five learning trials for 5XFAD mice. **(G)** Percentage of time spent in the inner zone in the NOR task and average speed (inches per second) for 5XFAD mice. **(H)** Input/output curves as a measure of basal synaptic transmission in the CA1 region of the hippocampus (n = 8-9 slices, 4-5 mice/group). **(I)** LC/MS measurements of KP metabolites in APP/PS1 and WT littermates with genetic deletion of *Ido1* (5-6 months old, n=6-7 mice /group). **(J)** Total Aß_42_ levels in cerebral cortex of vehicle or PF068-treated APP/PS1 mice. **(K)** Total Aß_42_ levels in cerebral cortex of APP/PS1 mice with genetic deletions of *Ido1*.

**Supplementary Fig. 6.**
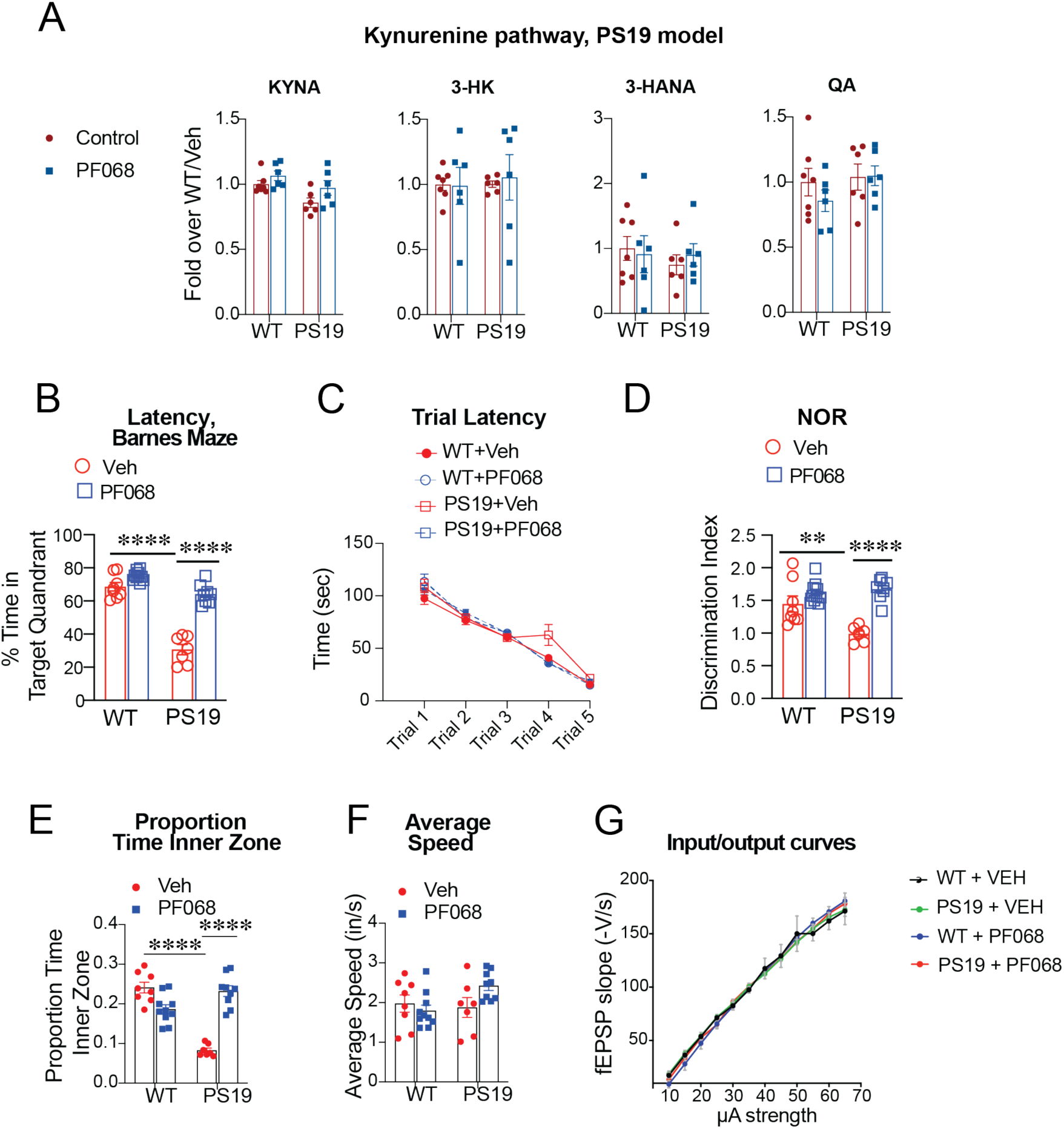
IDO1 inhibition in mutant tau PS19 mice. Data are mean ± s.e.m. and are analyzed using two-way ANOVA with Tukey post hoc tests: \**P*<0.05, \*\**P*<0.01, \*\*\**P*<0.001 and \*\*\*\**P*<0.0001. PS19 and WT littermates (8-9 months old) were administered 15 mg/kg/day PF068 or vehicle for 1 month and then tested. **(A)** LC-MS quantification of hippocampal KP metabolites (n=6/group). **(B)** Primary latency (sec) to the target hole in the Barnes Maze task (n=7-10/group). **(C)** Latencies in the Barnes maze for the five learning trials (n=7-10/group). **(D)** Discrimination index in the NOR task (n=7-10/group). **(E-F)** Percentage of time spent in the inner zone in the NOR task and average speed (inches per second; n=7-10/group) **(G)** Input/output curves as a measure of basal synaptic transmission in the CA1 region of the hippocampus in PS19 mice (n = 8-9 slices, 4-5 mice/group).

**Supplementary Fig. 7.**
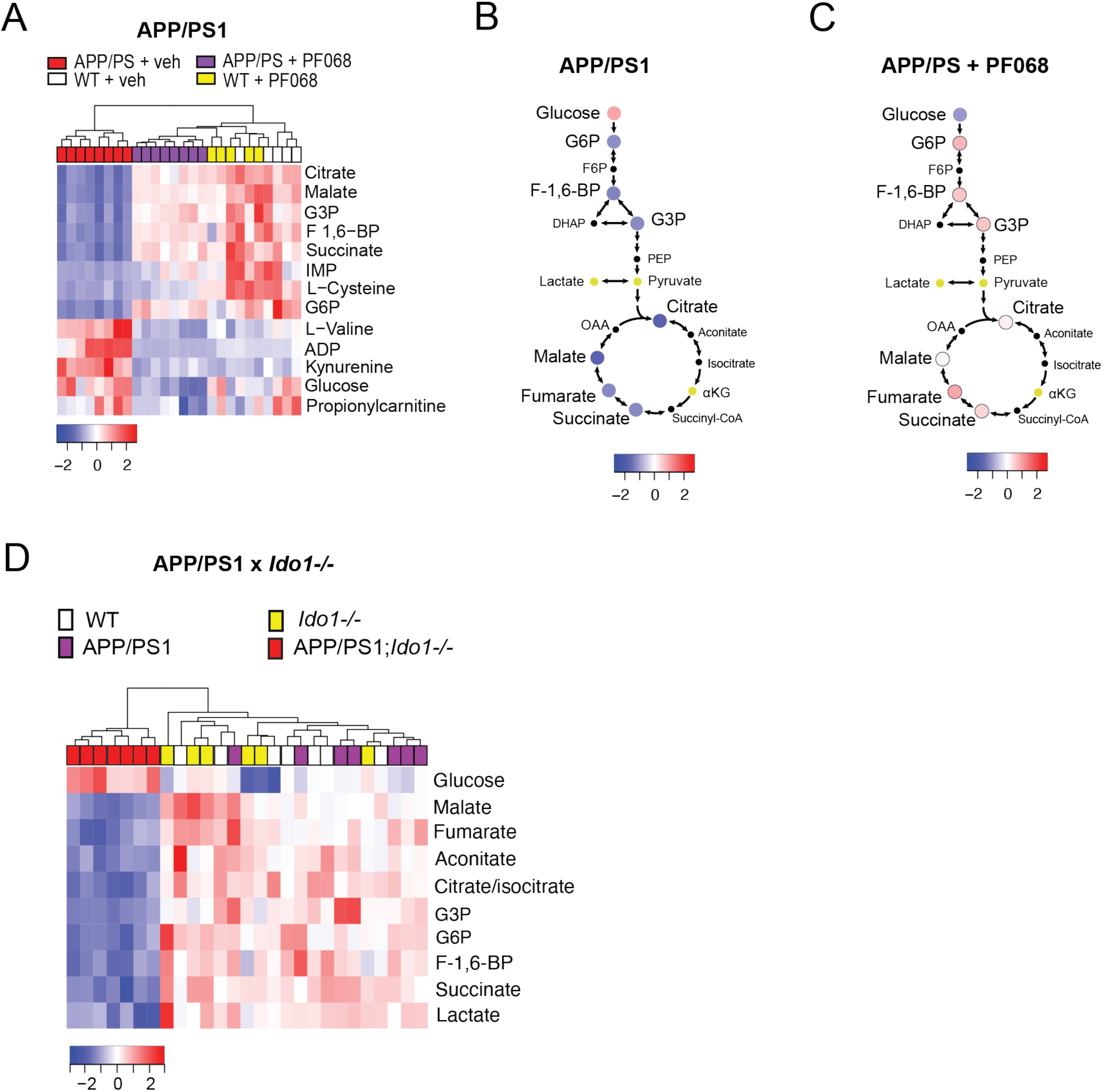
Pharmacologic or genetic inhibition of IDO1 improves glucose metabolism in hippocampus of APP/PS1 mice. **(A)** Hierarchical clustering of the 13 shared hippocampal metabolites in APP/PS1 and WT littermates +/- PF068 for one month (Z-score, -2 to 2). Hierarchical clustering is represented in terms of distance from the mean, or Z-score. **(B-C)** Schematic depicting levels of glycolysis and TCA metabolites and their average Z-score value from (**A**). Note the rescue of glycolysis downstream of glucose with IDO1 inhibition in APP/PS1 mice, similar to that observed in 5XFAD mice and PS19 mice. **(D)** Genetic knockout of *Ido1* also rescues glycolytic and TCA cycle metabolism in hippocampi of APP/PS1 mice. Glycolytic and TCA cycle metabolomic profiling of hippocampi isolated from WT (n=7), *Ido1-/-* (n=6), APP/PS1 (n=7), APP/PS1;*Ido1-/-* (n=7) littermates at 10-12 months of age. Note rescue of glycolytic metabolites, similar to pharmacologic model.

**Supplementary Fig. 8.**
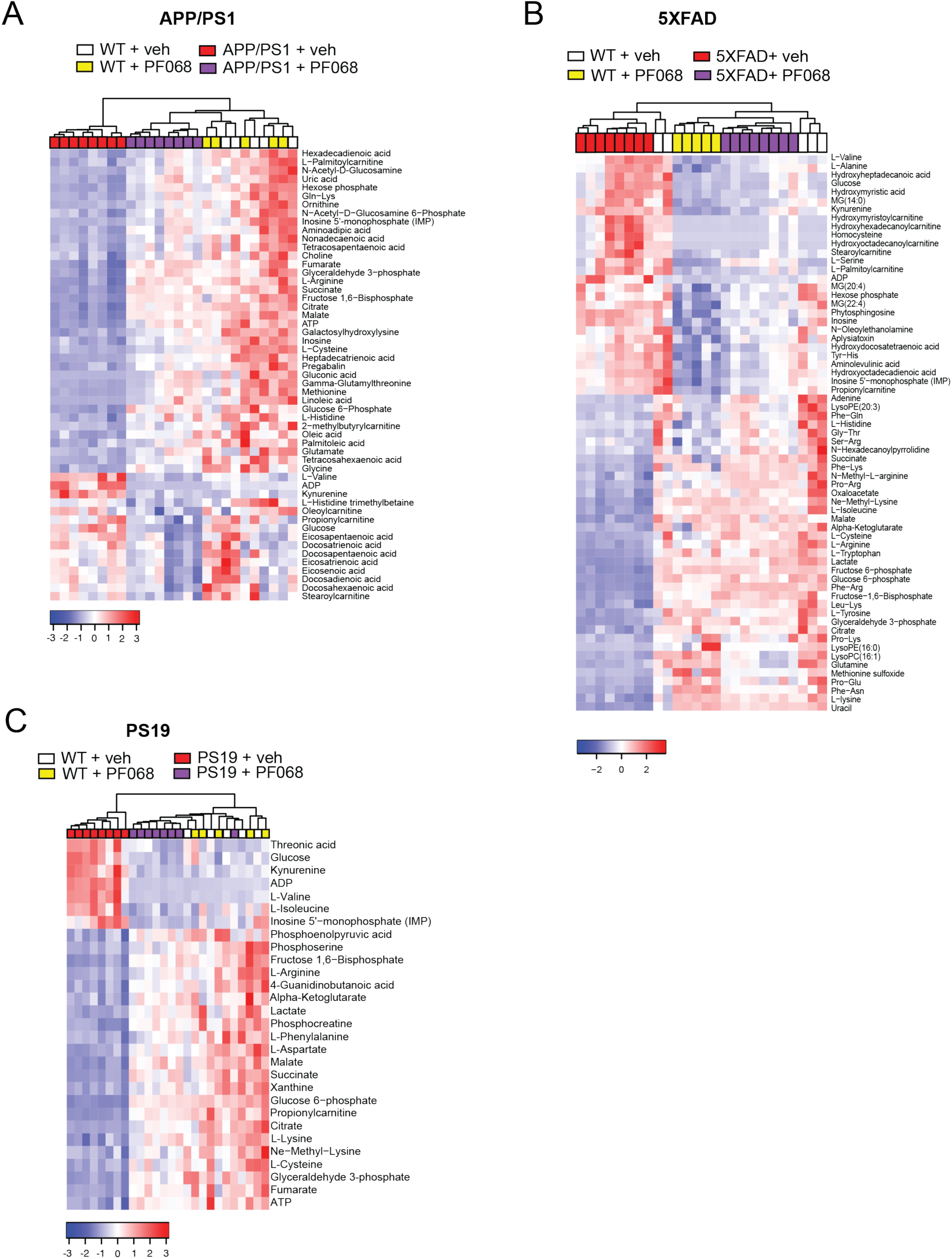
Metabolomic profiling of hippocampi from APP/PS1, 5XFAD, PS19 and WT mice +/- PF068 for 1 month. APP/PS mice (10-12 mo), 5XFAD mice (5-6 mo), and PS19 mice (8-9 months old) and wild type littermates were administered PF068 at 15 mg/kg/day for one month and then tested. Hierarchical clustering of all FDR-corrected metabolites is represented in terms of distance from the mean, or Z-score. (**A**) Hierarchical clustering of significantly altered metabolites (*q* < 0.05; APP/PS1 + PF068 vs. APP/PS1 + veh, n=5-8/group). (**B**) Hierarchical clustering of significantly altered metabolites (*q* < 0.05, 5xFAD + PF068 vs. 5xFAD + veh; n=5-8/group). (**C**) Hierarchical clustering of significantly altered metabolites (*q* < 0.05, PS19 + PF068 vs. PS19 + veh; n = 5-8/group).

**Supplementary Fig. 9.**
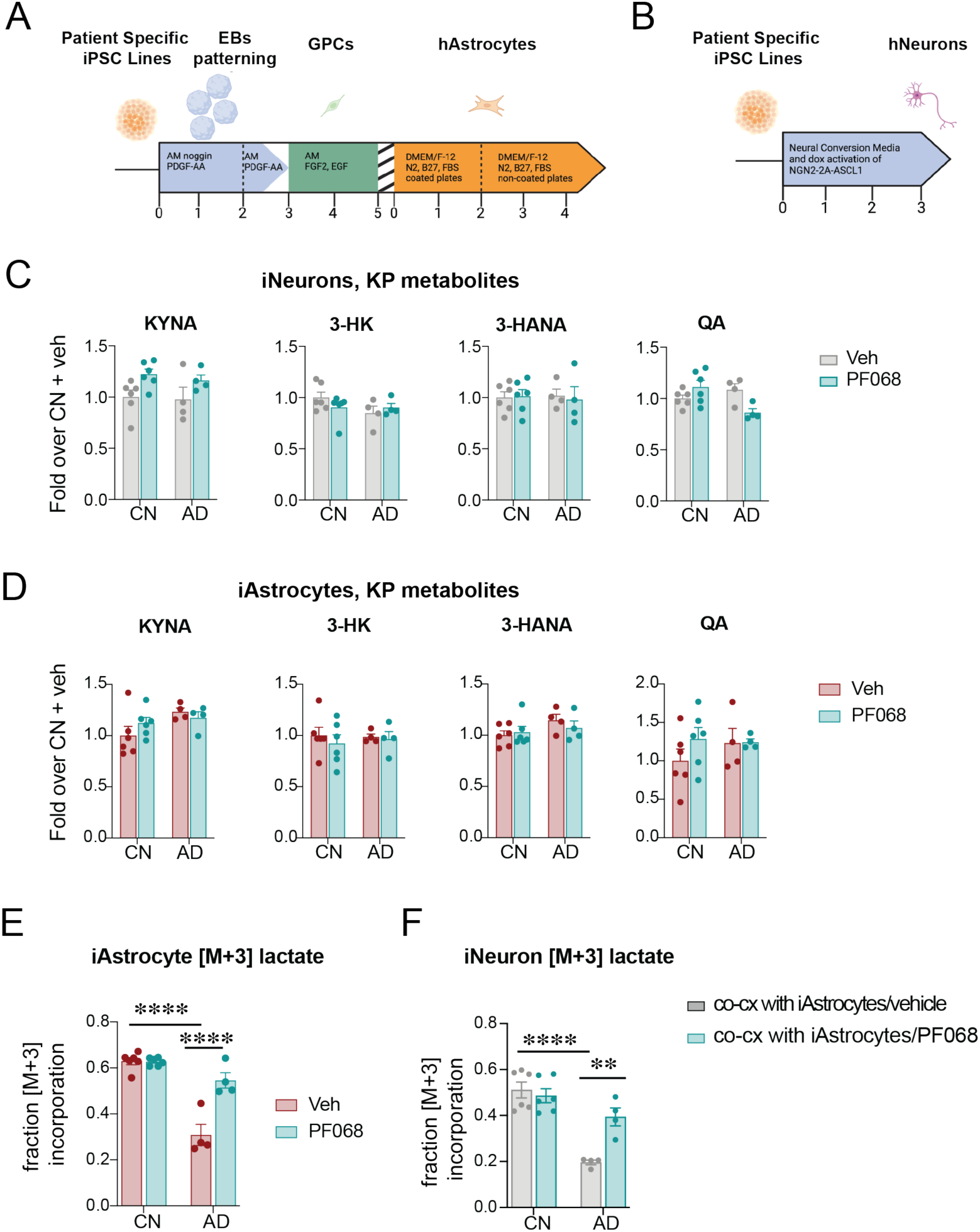
RNA-seq of hAstrocytes and hNeurons derived from CN and AD subjects. Data are mean ± s.e.m. and are analyzed using two-way ANOVA with Tukey post hoc tests: \*\**P*<0.01 and \*\*\*\**P*<0.0001. **(A)** Schematic of differentiation of hAstrocytes from subject specific iPSC line. **(B)** Schematic of differentiation of hNeurons from subject specific iPSC line. **(C)** KP metabolites downstream of KYN in hNeurons (n=6 control or CN, n=4 AD). **(D)** KP metabolites downstream of KYN in hAstrocytes (n=6 control or CN, n=4 AD). **(E)** Fraction of [M+3] lactate incorporated into hAstrocytes in response to IDO1 inhibition (n=6 control or CN, n=4 AD; PF 068 100 nM, 20 h). **(F)** Fraction of [M+3] lactate incorporated into hNeurons co-incubated with congenic hAstrocytes from CN or AD subjects +/- PF068 (n=6 control or CN, n=4 AD).

**Supplementary Fig. 10.**
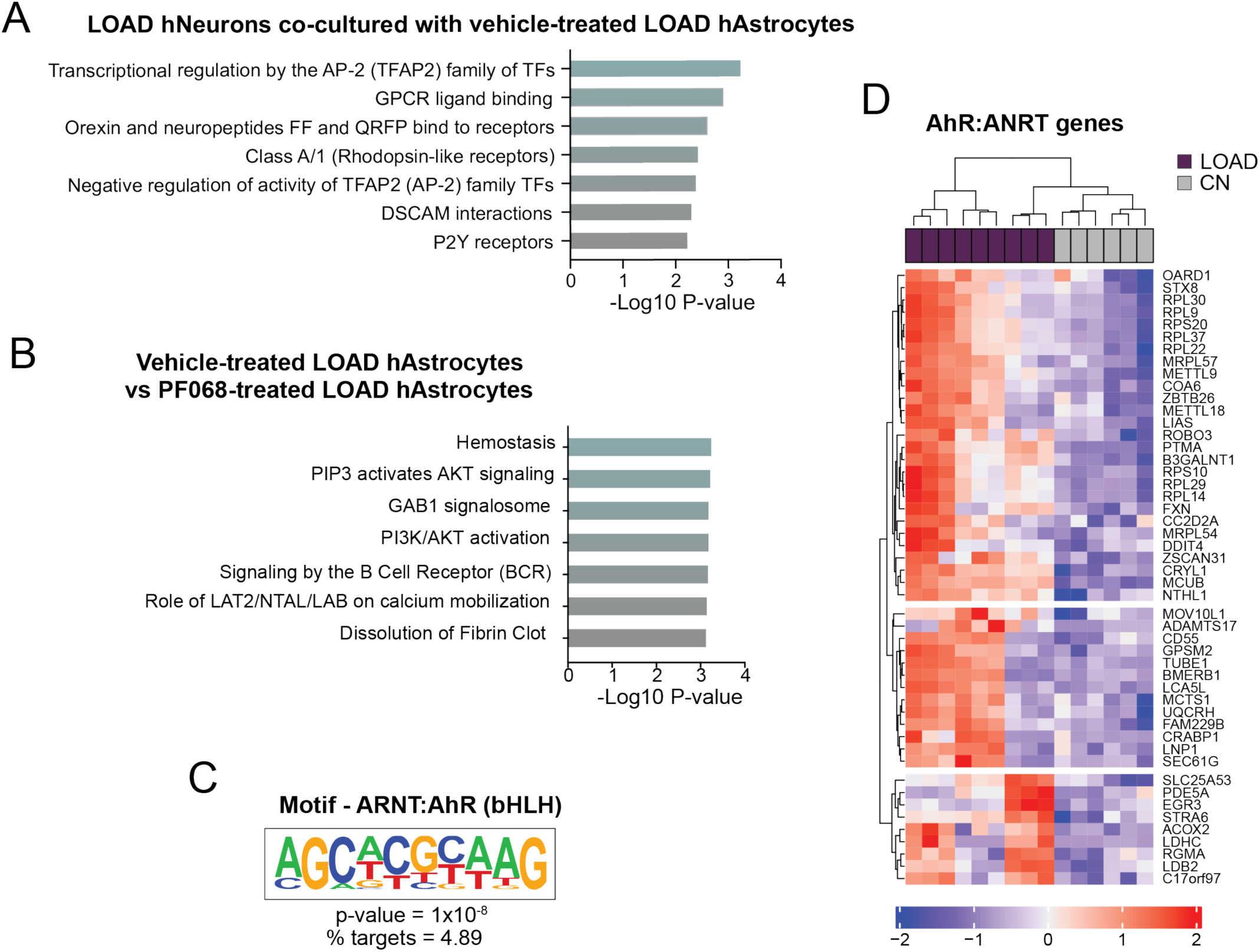
RNA-seq of co-cultured hNeurons and hAstrocytes from CN and LOAD subjects. **(A)** Top significantly enriched gene ontology terms from significantly upregulated transcripts in CN hNeurons co-cultured with hAstrocytes that were treated with vehicle. **(B)** Top significantly enriched gene ontology terms derived from significantly upregulated transcripts of hAstrocytes treated with vehicle versus the IDO inhibitor PF068. **(C)** Depiction of AhR/ARNT motif that is highly enriched in hAstrocyte RNA seq, n=4 AD and n=6 CN subjects. Homer was run for each biological comparison as well as four randomized gene sets to avoid false positive motif calls. **(D)** Hierarchical clustering and heatmap of transcripts identified by AhR motif analysis in hAstrocytes from n=4 AD and n=6 CN subjects.

